# Evolution of CDK1 paralog specializations in a lineage with fast developing planktonic embryos

**DOI:** 10.1101/817783

**Authors:** Xiaofei Ma, Jan Inge Øvrebø, Eric M Thompson

**Author notes:** Huntsman Cancer Institute, University of Utah, USA. **Abbreviations:** CDK, Cyclin Dependent Kinase; DoS, Determinant of Specificity; RNAi, RNA interference; NEBD, Nuclear Envelope Breakdown.

## Abstract

The active site of the essential, eukaryotic CDK1 kinase is generated by core structural elements, among which the PSTAIRE motif in the critical αC-helix, is universally conserved in metazoans. The CDK2 kinase, sharing the PSTAIRE, arose early in metazoan evolution and permitted subdivision of tasks along the S-M-phase axis. The marine chordate, *Oikopleura dioica*, is the only metazoan known to possess more than a single CDK1 ortholog, and all of its 5 paralogs show sequence divergences in the PSTAIRE. Through assessing CDK1 gene duplications in the appendicularian lineage, we show that the CDK1 activation loop substrate binding platform, ATP entrance site, hinge region, and main Cyclin binding interface, have all diversified under positive selection. Three of the 5 CDK1 paralogs are required for embryonic divisions and knockdown phenotypes illustrate further subdivision of functions along the S-M-phase axis. In parallel to CDK1 gene duplications, there has also been amplification in the Cyclin B complement. Among these, the CDK1d:Cyclin Ba pairing is required for oogenic meiosis and early embryogenesis and shows evidence of coevolution of an exclusive interaction. In an intriguing twist on the general rule that Cyclin B oscillations on a background of stable CDK1 levels regulate M-phase MPF activity, it is CDK1d protein levels that oscillate, rather than Cyclin Ba levels, to drive rapid, early embryonic cell cycles. Strikingly, the modified PSTAIRE of odCDK1d shows convergence over great evolutionary distance with plant CDKB, and in both *O. dioica*, and plants, these variants exhibit increased specialization to M-phase.

## Introduction

The complexity of the eukaryotic cell and its larger genome necessitates greater control over the organization of events during cellular division than for prokaryotic life. The eukaryotic protein kinase, CDK1, whose activity depends on unstable Cyclin regulatory subunits, has key roles in orderly triggering S and M phases from yeast to mammals (Santamaria et al. 2007), by sequential, regulatory phosphorylation of hundreds of substrate proteins. CDK activity levels progressively surpass response thresholds, to trigger serial events in S and M phases (Swaffer et al. 2016). Quantitative changes in CDK activity function as a coarse organizing mechanism for cell cycle transitions, with lower CDK activity resulting in higher affinity S-phase substrate phosphorylation, to carry out DNA replication, and higher activity driving mitotic entry via the phosphorylation of lower affinity mitotic substrates. CDK1 protein levels are constant during the cell cycle (Arooz et al. 2000), and are activated at each phase of the cell cycle by binding of stage-specific cyclins. The intrinsic activity of Cyclin–CDK1 complexes increases in correlation with the appearance of particular cyclins in the cell cycle (Ord and Loog 2019). G1/S cyclins generate CDK complexes with the lowest activity, which manifests in the lowest affinities (k_m_) and the lowest catalytic activities (k_cat_) toward substrates, whereas S-, G2-, and M-phase Cyclins form CDK complexes with higher affinities and catalytic activities towards substrates. The lower CDK activities generated by S- versus M-phase Cyclins also means that S-phase Cyclins do not achieve sufficient CDK1 activity levels to initiate mitosis, helping to ensure resolution of S- and M-phases. Compared to single-celled yeast, metazoans have also evolved CDK2, with lower intrinsic activity than CDK1 in the regulation of S–phase (Ord et al. 2019). Although metazoan CDK1 has become more specialized to M-phase functions it still retains indispensable, ancestral, late S-phase functions (Katsuno et al. 2009; Farrell et al. 2012; Seller and O’Farell 2018; Szmyd et al. 2019).

In addition to modulating CDK activity, the Cyclins are also implicated in specifying CDK substrate targets. Cyclins bind different substrate docking motifs: RxF for S Cyclins and LxF for budding yeast M Cyclins, and these Cyclin docking pockets increase specificity to target subsets to further fine-tune the CDK threshold ladder (Koivomagi et al. 2011; Ord et al. 2019). Most CDK substrates contain multiple target sites clustered in disordered regions (Holt et al. 2009), and phosphorylation cascades along these sites are shaped by precisely oriented docking interactions mediated by Cks1, the phospho-adaptor subunit of Cdk1 (Koivomagi et al. 2011). Distances between phosphorylation sites and docking sites are critical for both Cyclin and Cks docking. The composition of phosphorylation sites with respect to serine versus threonine, and surrounding residues, combined with positioning of Cyclin and Cks docking motifs, creates a unique barcode on each substrate. The Cyclin−CDK−Cks1 complexes read the barcodes and assign execution of CDK-triggered switches to specified time points during the cell cycle (Ord et al. 2019).

Canonical cell cycle regulation can be modified during development and growth. This is true of rapid mitotic cell cycles observed during early embryogenesis in a wide range of phyla and in the promotion of growth through endocycles in a number of organisms (Edgar and Orr-Weaver 2011).Externally developing, nutrient-rich, non-feeding, embryos are at risk of predation, and rapid development is a means of mitigating this risk (Strathmann et al. 2002). Many marine planktonic larvae also continue fast development to first swimming in order to enhance survival (Staver and Strathmann 2002). The pelagic, tunicate, *Oikopleura dioica*, exhibits very rapid embryonic and larval development, undergoing metamorphosis to a feeding juvenile within 12 h at 15 °C (Bouquet et al. 2009). Very rapid growth and extensive modulation of reproductive output over several orders of magnitude are then achieved through deployment of a variety of endoreduplicative cell cycle variants (Ganot and Thompson 2002; Ganot et al. 2007). These adaptations allow the organism to rapidly adjust population levels in response to algal blooms, during an extremely short chordate life cycle (6 days at 15 °C). Accompanying this life history strategy is considerable modification of the organism’s molecular cell cycle regulatory complement (Campsteijin et al. 2012), including gene duplications to generate multiple CDK1 and Cyclin B paralogs.

All metazoans examined to date possess a single CDK1 ortholog that exhibits a highly conserved PSTAIRE motif in the critical αC-helix structural element located within the N-lobe of CDK1 and CDK2. This element is an important interface in docking interactions with Cyclin partners and in contributing to formation of the kinase active site (Wood et al. 2019). *O. dioica*, aligns with other metazoans in possessing a CDK2 with a conserved PSTAIRE motif, but deviates distinctively in that it has evolved 5 CDK1 paralogs, all of which show deviations in the critical PSTAIRE motif. Here, we have examined how these paralogs have arisen in the appendicularian lineage and to what extent they exhibit functional specializations along the CDK2-CDK1, S- to M-phase, cell cycle, regulatory axis. We show that the CDK1 substrate binding platform, activation loop, ATP entrance site, hinge region, and the main Cyclin binding interface, have all diversified under positive selection. In meiosis and early embryogenesis, CDK1 paralogs were sequentially activated across the cell cycle. Combinatorial knockdowns revealed collaboration of CDK2, CDK1a, CDK1b and CDK1d along the S- to M-phase axis in oogenic meiosis and early embryonic divisions. Interestingly, the Cyclin Ba and CDK1d paralogs are located in proximity on the recently evolved *O. dioica* X chromosome, were co-expressed specifically during oogenic meiosis and early embryogenesis, and were required for these processes. Modifications of critical residues in the Cyclin B:CDK1 interface and the salt bridge, suggest that CDK1d and Cyclin Ba may have coevolved an exclusive interaction that does not interfere with the more canonical CDK1a/b/c:CycBc interactions in regulating M-phases during early developmental events. It is generally universal in metazoan mitotic cycles that CDK1 protein levels remain relatively constant throughout the cell cycle whereas Cyclin B levels oscillate and are up-regulated during M-phase. In contrast, we reveal a novel regulatory oscillation of levels of the CDK1d paralog, as opposed to its Cyclin Ba interacting partner, that drive early, very rapid embryonic cell cycles in this planktonic chordate.

## Results

### Episodic positive selection of CDK1 orthologs after duplication

Metazoans possess a single CDK1, with the exception of *Oikopleura dioica*, where 5 CDK1 paralogs have been identified (Campstejin et al. 2012). To trace the origins of this amplification we analyzed genomes and transcriptomes of five appendicularian species in the *Oikopleura* and *Fritillaria* genera (Naville et al. 2019). *CDK1a, b and c* were uniformly present, whereas *CDK1d* and *e* appear to have arisen from *CDK1c* specifically in *O. dioica*, the only known dioecious appendicularian (fig. 1A). Interspecies clustering of *CDK1* paralogs and interspecies conservation of their intron-exon structures (supplementary fig. S1) suggest that *CDK1* amplification arose through early DNA-based duplication events in the *Oikopleura* lineage. Based on phylogenetic analyses, *CDK1a* duplication gave rise to *CDK1b* that was subsequently duplicated to generate *CDK1c*, which then gave rise to the *O. dioica* specific, *odCDK1d*. This duplication gradient pattern, and young paralogs-biased duplication, have not previously been reported in metazoans. *O. albicans CDK1a*, *b* and *c* locate on the same chromosome. *O. dioica CDK1a* and *b* locate on the same autosome whereas *CDK1c* is present on a different autosome that also contains *CDK1e* (fig. 1C). *CDK1d* is found on the X chromosome where, at a distance of 5 MB, the *CycBa* locus is also located. Following the classification system for paralog subtypes (Sonnhammer and Koonin 2002), *Oikopleura* genes in the *CDK1a* clade are orthologs, and *odCDK1a* and *b* are outparalogs to each other when comparing *O. dioica* with *O. albicans*, since CDK1 duplication happened before *O. dioica* and *O. albicans* speciation. *O. dioica CDK1c*, *d* and *e* are inparalogs to each other when comparing *O. dioica* with *O. albicans*, since CDK1c duplication happened after *O. dioica* and *O. albicans* speciation.

**FIG. 1.**
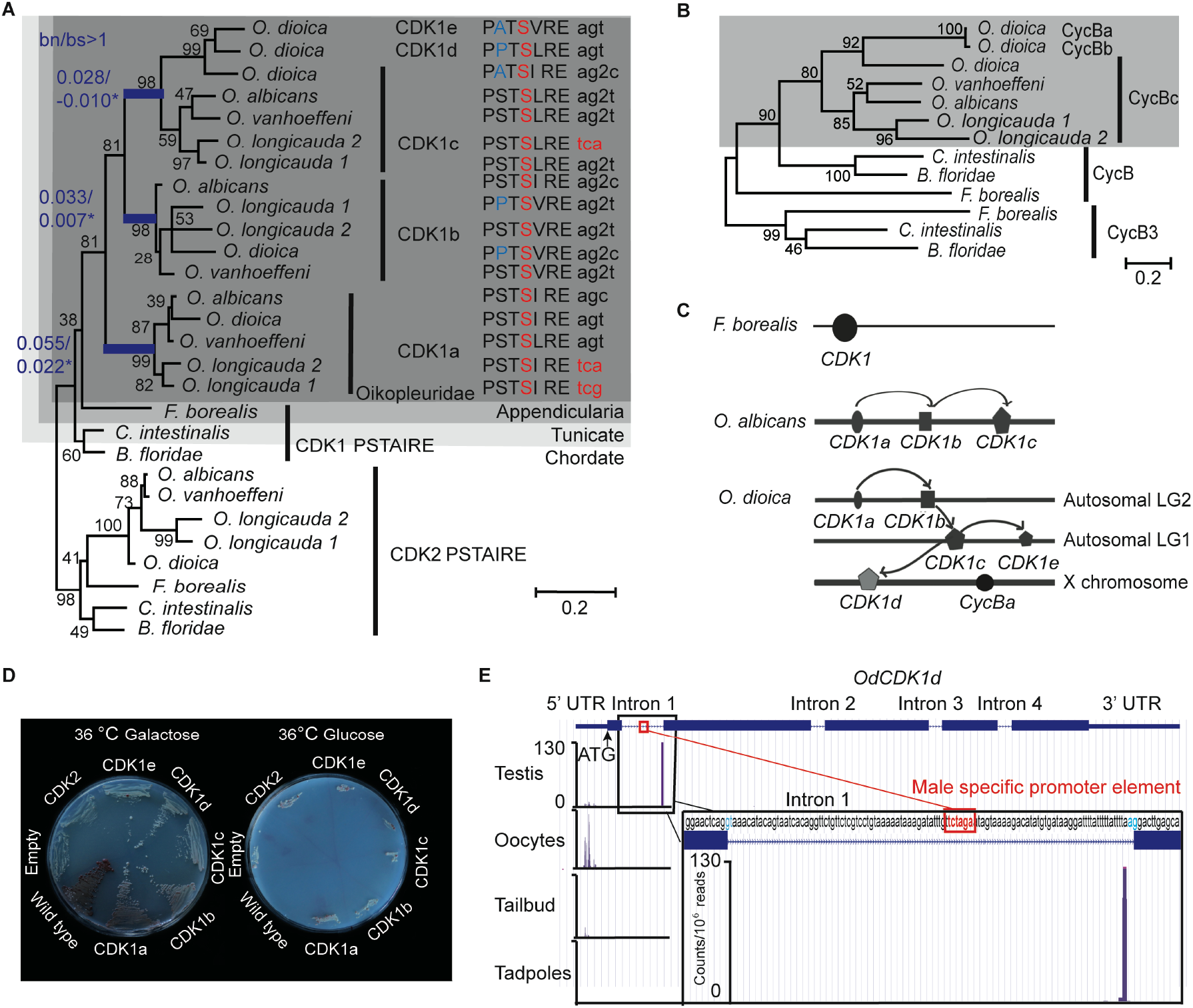
Amplification of CDK1 paralogs in the *Oikopleura* genus. (*A*) Left panel: Maximum likelihood inference analysis of selected chordate CDK1 proteins using the kinase domain of CDK1s with bootstrap values indicated at nodes. bn/bs values in all branches were calculated using a modified Nei-Gojobori method (Zhang et al. 1998). Blue branches have significant bn/bs values over 1 as determined by one-sided Z tests: lineage subtending CDK1a, P = 0.0386; lineage subtending CDK1b, P = 0.0305, lineage subtending CDK1c, P = 0.0334. Modifications in the highly conserved PSTAIRE domain of each CDK1 are shown. The codons resulting in the alanine to serine substitution within the Oikopleuridae (dark grey box) are also indicated. Codons for serine with phase 2 intron insertions (splitting codons between the second and third nucleotides) are denoted by “2”. (*B*) Maximum likelihood inference analysis of selected chordate Cyclin B proteins. The Oikopleuridae paralogs are indicated by a grey box. (*C*) CDK1 paralog localizations on chromosomes among appendicularians. Arrows indicate possible chronological duplication steps and chromosomal translocations consistent with the phylogenetic analyses. (*D*) Complementation studies using budding yeast cdc28-4ts mutants substituted with *O. dioica* CDK1 paralogs. CDK1a, b and d paralogs were able to complement whereas c and e did not. (*E*) Alternative *CDK1d* transcription start sites (TSS) in testes versus ovaries. CDK1d maternal transcripts have a broad TSS before the ATG start codon. Testes *CDK1d* transcripts originated from a single TSS immediately before the AG splice acceptor site of fist intron, 42 nt downstream of the TCTAGA male-specific promoter element, which is also located within the first intron. Male *CDK1d* transcripts lack an ATG start codon, and yield nonsense transcripts in all three possible reading frames. Vertical axis scale for CAGE data (Danks et al. 2018) is in reads per million.

Both the new Nei-Gojobori method (Zhang et al. 1998) and Codon-based models (Yang 2007) were applied to analysis of *Oikopleura* CDK1s. During the 3 rounds of gene duplication, the lineage subtending *CDK1a* (fig. 1A) experienced positive selection (bn/bs = 2.48, P=0.0386, dN/dS(ω) = 41.01, p = 0.081, P = 0.0481), as did *CDK1b* (bn/bs = 4.72, P = 0.0305, dN/dS(ω) = 28.14, p = 0.046, P = 0.003) and *CDK1c* (bn/bs = infinite, P = 0.033, dN/dS(ω) = 999, p = 0.066, P=0.002) (fig. 1A, supplementary table S1). To better understand positively selected sites identified by Codon-based models, we aligned CDK1 sequences from 15 metazoan phyla, and Cdc28 from budding yeast. Sites conserved in metazoan sequences, but substituted in *Oikopleura* CDK1 orthologs, were mapped to human CDK1 and CDK2 crystal structures (supplementary figs. S2 and S3). Determinants of kinase specificities (DoS) (Creixell et al. 2015) were aligned and the DoS that are conserved in metazoan CDK1 orthologs but substituted in *Oikopleura* CDK1 sequences were mapped to human CDK1-Cyclin B1 crystal structures. During kinase evolution, the critical catalytic residues, regulatory hydrophobic spine and catalytic hydrophobic spine have been conserved. The diversification of the substrate binding platform-activation segment, and indirect ATP binding sites, drive diversification of substrate specificity and substrate phosphorylation efficiency. *Oikopleura* CDK1s have the same active site as human *CDK1* and yeast cdc28, including the Mg^2+^-binding loop (DFG), important residues in the catalytic loop, the catalytic hydrophobic spine, the regulatory hydrophobic spine and the salt bridge (Wood et al. 2019) (supplementary figs. S2 and S3). The bn/bs values below 1, observed among CDK1 sequences (fig. 1A), suggest that these orthologs are functional kinases under purifying selection. Conservation in the DFG+1 residues (Leu) in the 17 *Oikopleura* CDK1 sequences, suggests that they share with human CDK1, a preference for serine over threonine as a phosphor-acceptor residue in substrates (Chen et al. 2014). However, the ATP binding sites, substrate binding sites and Cyclin binding interfaces have diversified in *Oikopleura* CDK1 paralogs.

### Substitutions in ATP binding sites

In the protein kinase family, the ATP binding site is very conserved, but adjacent areas not occupied by ATP are more variable, particularly in the hinge region (E81-K89, human CDK1 numbering). These sites modulate ATP affinity. This region may determine the different catalytic efficiencies of CDK1 and CDK2, as substitutions in the hinge of CDK2, that mimic amino acids in this region of CDK1 (substitute N84, Q85 in CDK2 to 84S, 85M in CDK1), increase both ATP affinity (k_m_ ATP) and turn over (k_cat_) (Echalier et al. 2012). In the opposite direction, a K89D/E mutation in the CDK2 sequence, reduces k_cat_. *Oikopleura* CDK1 paralogs have diversified in the hinge region. CDK1c paralogs exhibit an S84F/Y substitution whereas this site is conserved in CDK1a paralogs compared to the metazoan consensus (supplementary fig. S2). The M85 site is a DoS, and a M85C substitution occurred in CDK1a of *O. vanhoeffeni*, *O. albicans* and *O. dioica*, after they split from *O. longicauda.* This substitution has been positively selected (supplementary table S1), possibly driving diversification of ATP affinity among CDK1 paralogs following their duplication. *Oikopleura* CDK1a and b orthologs have substitutions at K89, a residue that is conserved in CDK1 and CDK2 from yeast to metazoans (supplementary fig. S2). This raises the possibility that CDK1a and b may have lower ATP affinity and substrate phosphorylation efficiency compared to CDK1c, d and e paralogs. Indeed, as shown later, we found that CDK1d peaks in M phase whereas CDK1a was present earlier in interphase. The CDK1 L83 residue is also conserved form yeast to human, with the amide of L83 forming a hydrogen bond with the heteroaromatic adenine ring of ATP (Bao et al. 2011). This residue is conserved in CDK1a and c, but has been modified to L83M in the hinge region of all *Oikopleura* CDK1b orthologs. Thus, CDK1 paralogs have different signatures in their hinge regions, suggesting diversification in their ATP affinities and catalytic efficiencies.

The conserved DoS, P61, is located in the N-lobe, in the loop connecting the PSTAIRE helix and β4 of human CDK1 (supplementary fig. S2). Positive selection of a P61E substitution in all *Oikopleura* CDK1a orthologs, suggests that positive selection occurred after the initial CDK1 duplication but prior to speciation. This asymmetric adaptive evolution of an evolutionary conserved DoS in one of the duplicates, and conservation of this substitution within CDK1a orthologs, could be best explained by escape from adaptive conflicts after duplication (Des Marais and Rausher 2008).

### Substitution in the activation segment

The activation segment forms an integral part of the substrate binding groove, and diversity in this loop contributes to the wide range of substrate specificities and affinities of protein kinases (Nolen et al. 2004). The structure of CDK1 suggests a mechanism through which activation loop flexibility, embedded in a more flexible CDK1 fold, allows CDK1 to accommodate a more diverse substrate set than CDK2 (Brown et al. 2015). These plastic properties may also contribute to its ability to partner non-cognate cyclins in the absence of other CDKs to drive a complete metazoan cell cycle^1^. In *Oikopleura* CDK1 paralogs, sites in the activation segment that bind to substrate, P0 (L149, T166), P+1 (V164, V165) and P+3 (T161), are the same as in yeast and human CDK1 (supplementary fig. S2). However, several conserved sites in the activation segment have been dramatically modified in *Oikopleura CDK1* paralogs, particularly in *O. dioica CDK1c, d* and *e* inparalogs. G154 is located at the tip of the human CDK1 activation segment where it is important in forming the β turn in the short β hairpin structure when CDK1 binds to Cyclin B1 (Brown et al. 2015) (supplementary fig. S3). Human CDK2 cannot form this hairpin structure (Jeffrey et al. 1995), and this is a major difference in the activation segment between CDK1 and CDK2. The G154 site is conserved in yeast and metazoans, but has diversified in *Oikopleura* CDK1 paralogs. A G154N substitution occurred in *Oikopleura* CDK1a, G154S in *Oikopleura* CDK1b, and G154K in *O. dioica* CDK1c, d and e (supplementary fig. S2). This site is also identified as having experienced positive selection (supplementary table S1), suggesting adaptive diversification of the activation segment by disrupting the β-turn though substitution with large side chain residues. Immediately downstream of this substitution, *O. dioica* CDK1c, d and e all have an I157F substitution, whereas CDK1c paralogs from other *Oikopleura* species have an I157M substitution. *O. dioica* CDK1c, d and e also exhibit substitutions at theV159 and Y160 sites, two residues immediately upstream of the important, regulatory phospho T161 site. The V159 site is also a DoS, conserved from yeast to humans. Finally, in human CDK1, L167 (conserved from yeast to human) is adjacent to the substrate P-2 residue (Brown et al. 1999; Bao et al. 2011) (supplementary fig. S4). The L167M substitution found in all CDK1b and c orthologs, presents a larger side chain, which may affect affinity towards P-2 residues.

### Substitutions in the PSTAIRE helix

The CDK1 PSTAIRE helix is invariant in metazoans and yeast. Mutation screens in yeast *cdc28* revealed that the exact PSTAIRE helix sequence (PSTAIREISLLKE) is required for cell division, and any mutations in this domain are lethal (MacNeill and Nurse 1993; Levine et al. 1999; Ahn et al. 2001;). The A48V substitution in cdc2 in *S. pombe* is dominant negative, as the mutant cdc2 could bind Cyclin but could not be activated (Fleig et al. 1992), whereas an A48T substitution in *S. cerevisiae cdc28* inhibits CLB2 association (Ahn et al. 2001). Surprisingly, the PSTAIRE helix is modified in all *Oikopleura* CDK1 paralogs (fig. 1A) whereas *Oikopleura* CDK2s retain the canonical PSTAIRE sequence. All of the 17 identified *Oikopleura* CDK1 sequences share the A48S substitution (supplementary figs. S1 and S2). The S48 residues in *Oikopleura* CDK1 sequences are encoded by two disjoint codon sets, TCN and AGY, which cannot be interconverted by a single nucleotide mutation. Accordingly, switches between serine codons from the two sets can occur either directly, by simultaneous double (tandem) mutation (TC > AG), or indirectly, via two consecutive single-nucleotide substitutions (TC > AC > AG or TC > TG > AG). This is consistent with strong positive selection (Rogozin et al. 2016) at these sites in CDK1 sequences within the entire *Oikopleura* genus. Interestingly, coincident with the A48S substitution, codons for S48 in all CDK1b and c paralogs, except olCDK1c, have phase 2 intron insertions (supplementary fig. S1). This intron insertion is specific to *Oikopleura*, and is not found in any other metazoan CDK1.

*O. dioica* CDK1b and d (and *O. longicauda* CDK1b) have an additional S46P substitution in the PSTAIRE helix whereas *O. dioica* CDK1c and e have an S46A substitution at this residue. The double proline predicts that the odCDK1b and d PSTAIRE helices would be shorter and exhibit an altered orientation compared to human CDK1. The *O. dioica* CDK1a, b and d versions of the PSTAIRE helix complemented yeast *cdc28* temperature sensitive mutants, whereas *O. dioica* CDK1c and e did not (fig. 1D). *O. dioica* CDK1c, d and e inparalogs also share double S53C and L54T/S substitutions in the PSTAIRE helix (supplementary fig. S2). Human CDK1 S53 forms hydrogen bonds with Cyclin B1 L296 and G297 amide groups (Brown et al. 2015) (located at the C-terminal loop connected to the α5 helix) (supplementary fig. S3). Substituting serine at this site with cysteine may reduce CDK1c inparalog interaction interfaces with Cyclin B.

In summary, critical elements for catalysis are conserved in *Oikopleura* CDK1 paralogs, but they diverge in binding sites for ATP and substrates, suggesting qualitative differences in catalysis. Therefore, we set out to determine expression profiles and evaluate knockdowns of *O. dioica* CDK1 paralogs to characterize their individual functions in meiosis and early embryonic cell cycles.

### Transcription of *CDK1d* and *cycBa* is restricted to oogenesis

*CDK1a, b, c* and *cycBc* are generally present in the *Oikopleura* genus (fig. 1A and B). In *O. dioica*, they were broadly expressed, in a sexually unbiased manner, in embryos, juveniles, and mature animals (fig. 2). In contrast, *CDK1d* and *e*, and *cycBa* are thus far, specific to *O. dioica*, and were transcribed during oogenesis. Their transcripts persisted during early embryogenesis (figs. 2 and 6B). *CycBa* was not expressed in juveniles or testes. Interestingly, *O. dioica CDK1d* has different transcription start sites (TSS) in testes and ovaries. The ovarian *CDK1d* TSS occurs before the ATG start codon located in the first exon. In testis, a sharp TSS located within the first intron yields nonsense transcripts (fig. 1E). Western blots confirmed that the CDK1d protein was present in ovaries but absent in testes (fig. 2B). *CDK1d* mRNAs began to degrade after fertilization and dropped to very low levels at 2.5 h post-fertilization (fig. 6B). Given these differential expression patterns, we then undertook to knockdown the five CDK1 paralogs and 2 Cyclin Bs in *O. dioica* individually, or simultaneously, to explore functions of the expanded CDK1-Cyclin B complement.

**FIG. 2.**
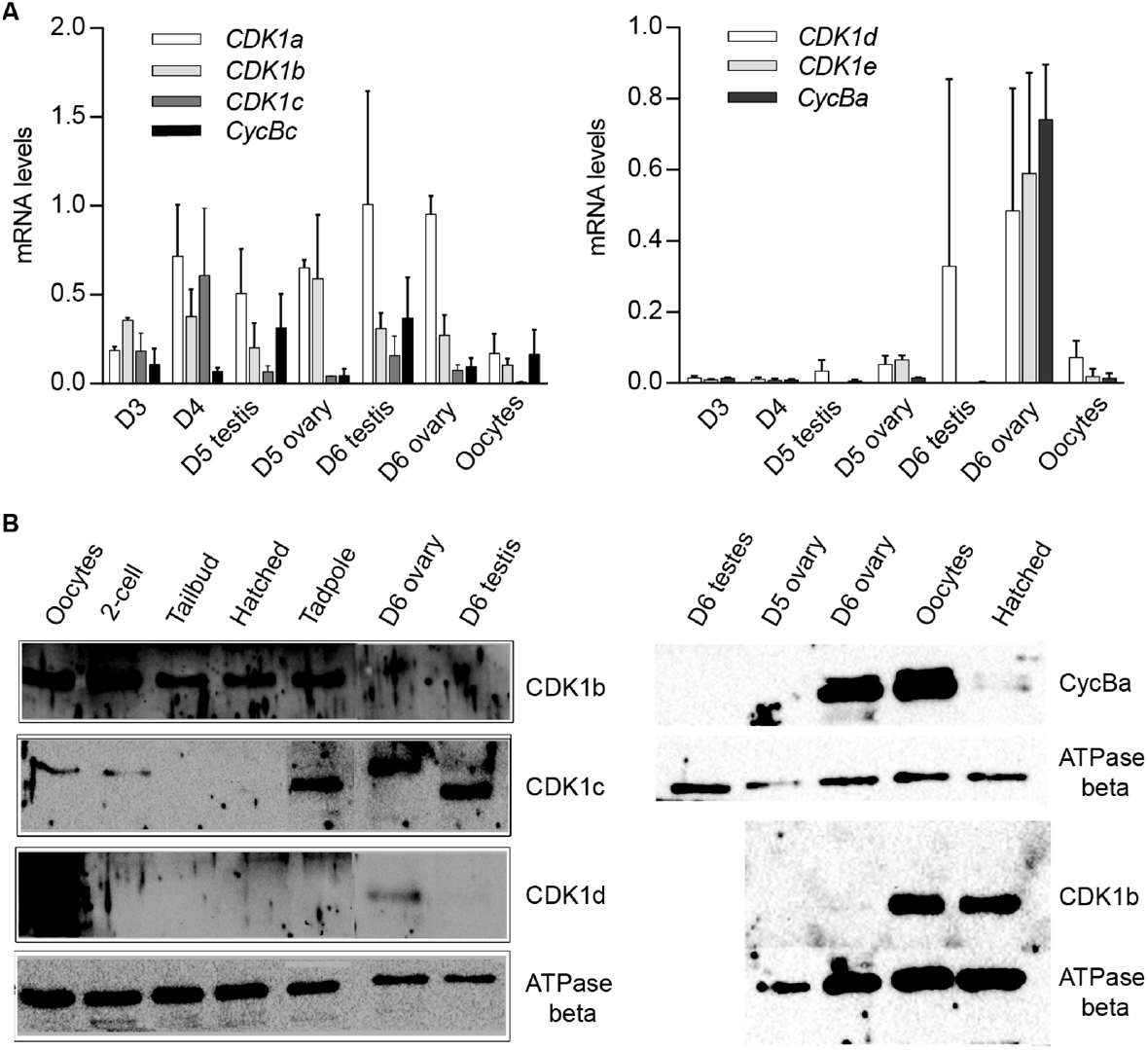
CDK1 and Cyclin B paralog expression profiles. (*A*) *CDK1-Cyclin B* paralog expression levels during gonad maturation from Day 3 to spawning. Expression data were normalized to transcription of the EF1b and Rpl23 housekeeping genes. Levels are normalized to maximum levels. (*B*) Western blots of CDK1-Cyclin B paralog levels during development, gonad maturation and embryogenesis.

### CDK1a, b and d collaborate in embryonic divisions: a conserved PSTAIRE is not required

The five odCDK1 paralogs and odCDK2 were each targeted for knockdown by RNAi. After injections of dsRNA in gonads of day 4 or 5 animals, oocytes were collected and processed to verify specific knockdown by RT-qPCR, and western blots. Successfully knocked down oocytes were exposed to wild type sperm and embryonic divisions were examined. Previous reports of CDK1a RNAi at day 4 demonstrated that this impaired vitellogenesis and meiosis resumption (Ovrebo et al. 2015; Feng and Thompson, 2018). CDK1a RNAi at day 5 avoided these defects and injected animals produced normal size oocytes. However, embryonic divisions were delayed and they failed to hatch (fig. 3A). When CDK1b was knocked down, embryonic cell cycles were also delayed and embryos arrested at gastrulation, corresponding to 2.5 h post-fertilization of normal embryos (fig. 3B). Knockdown of CDK1d yielded 3 phenotypes: a complete failure to emit polar bodies, arrest after polar body emission, or abnormal divisions after polar body emission (fig. 3C-G). These phenotypes reflect roles of odCDK1d in nuclear envelope breakdown (NEBD) (Ovrebo et al. 2015) and spindle assembly in meiosis I, for meiosis completion (Feng and Thompson 2018), and subsequent mitotic divisions. No significant differences were observed in the proportions of normal hatching embryos between wild type, and any of the *CDK1c, e*, or *c*+*e* RNAi knockdown embryos (supplementary fig. S5). CDK1c and e have very similar PSTAIRE helices (PATSI/VRE, fig. 1A), they failed to rescue yeast *cdc28* mutants, and they were also dispensable for embryonic divisions.

**FIG. 3.**
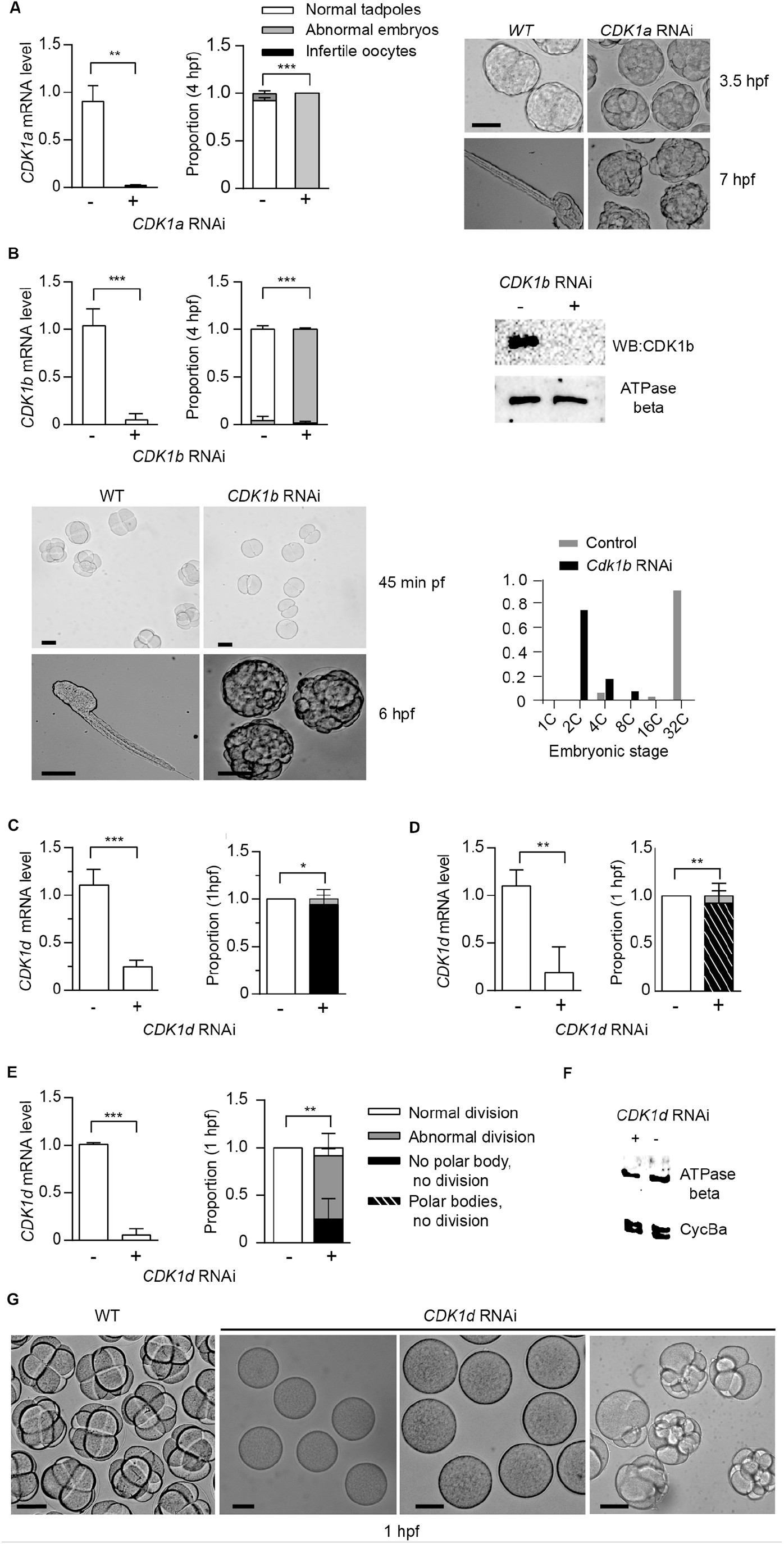
*CDK1a*,*b*, and *d* knockdowns generate abnormal embryonic phenotypes. (*A*) *CDK1a* RNAi delays embryonic divisions and tadpoles fail to form. Left panel: efficient knockdown of *CDK1a* in oocytes spawned from ovaries injected with dsRNA against *odCDK1a* at D5. Mid panel: When exposed to wild-type sperm, *CDK1a* deficient oocytes generated embryos that divided more slowly than wild-type and failed to hatch at 4 hpf. Right panel: images of delayed development in CDK1a deficient embryos. When wild-type embryos had developed to the late tailbud stage, CDK1a deficient embryos had only begun to gastrulate. At 7 hpf, tadpoles were observed in wild-type whereas CDK1a deficient embryos failed to reach the tailbud stage. The legend in (*A*) also applies to the respective panel in (*B*). (*B*) Upper left panel: efficient knockdown *CDK1b* in oocytes spawned from ovaries injected with dsRNA against *odCDK1b* at D5. Upper mid panel: When exposed to wild-type sperm, *CDK1b* deficient oocytes generated embryos that failed to hatch at 4 hpf. Upper right panel: Western blot showing absence of CDK1b protein in *CDK1b* RNAi embryos. Bottom left panel: CDK1b deficient embryos arrested before gastrulation. Lower right panel: distribution of developmental stages in *CDK1b* deficient and wild type embryos assessed at 1 hpf. (*C-E*), Knockdown of *CDK1d* generated 3 different phenotypes at 1 hpf. Left panels reveal efficient knockdown in oocytes spawned from ovaries that had been injected with dsRNA against *odCDK1d* at D5. Right panels revealed abnormal division in all cases with a subset of embryos exhibiting more severe phenotypes of polar body extrusion but no cell division, or a complete lack of polar body extrusion and absence of any division. The legend in (*E*) also applies to the respective panels in (*C*) and (*D*). (*F*) Knockdown of CDK1d did not have any effect on the levels of Cyclin Ba, the Cyclin component of MPF in *O. dioica* oocytes (Feng and Thompson 2018). (*G*) Image comparisons of wild-type to *CDK1d* knockdown oocyte/embryo phenotypes 1 h post exposure to wide type sperm. Left to right: WT cleavage stage embryos, infertile oocytes with no polar body extrusion, polar bodies extruded, but no division, and abnormally dividing embryos. Where indicated, ***P<0.001, **P<0.01, *P<0.05. Scale bars: 50 μm.

CDK2 is dispensable for early mouse early embryogenesis and mitotic cycles in cultured vertebrate cells (Santamaria et al. 2007). Consistent with these observations, *odCDK2* RNAi embryos divided normally and hatched with correct timing (fig. 4A). CDK2 is the only CDK in *Oikopleura* that has a canonical PSTAIRE helix. This indicates that the exact sequence in the conserved metazoan CDK1, PSTAIRE helix, is not required for *Oikopleura* embryonic divisions. After emission of polar bodies, *CDK1a* + *CDK2* RNAi zygotes arrested prior to pronuclear fusion (fig. 4B). The synergistic defects during the cell cycle caused by *CDK1a*+*CDK2* RNAi suggests CDK1 and CDK2 have overlapping roles in interphase. Since *CDK2* RNAi embryos divided normally, defects arising from *CDK1a*+*CDK2* RNAi suggest *CDK1a* could complement CDK2’s roles in interphase during embryonic divisions. Immunostaining of tadpoles developed from oocytes injected with *CDK1a-GFP* capped mRNA, showed CDK1a located to the nucleus during interphase (supplementary fig. S6). *CDK1a*+*b* RNAi embryos arrested at the 4^th^ or 5^th^ division at prophase with super-condensed chromosomes closely juxtaposed to the nuclear lamina (fig. 4D). The synergistic defects during the cell cycle cause by *CDK1a*+*b* RNAi suggests collaborative roles of CDK1a and CDK1b during prophase, the point at which CDK1b localized to chromatin (supplementary fig. S6).

**FIG. 4.**
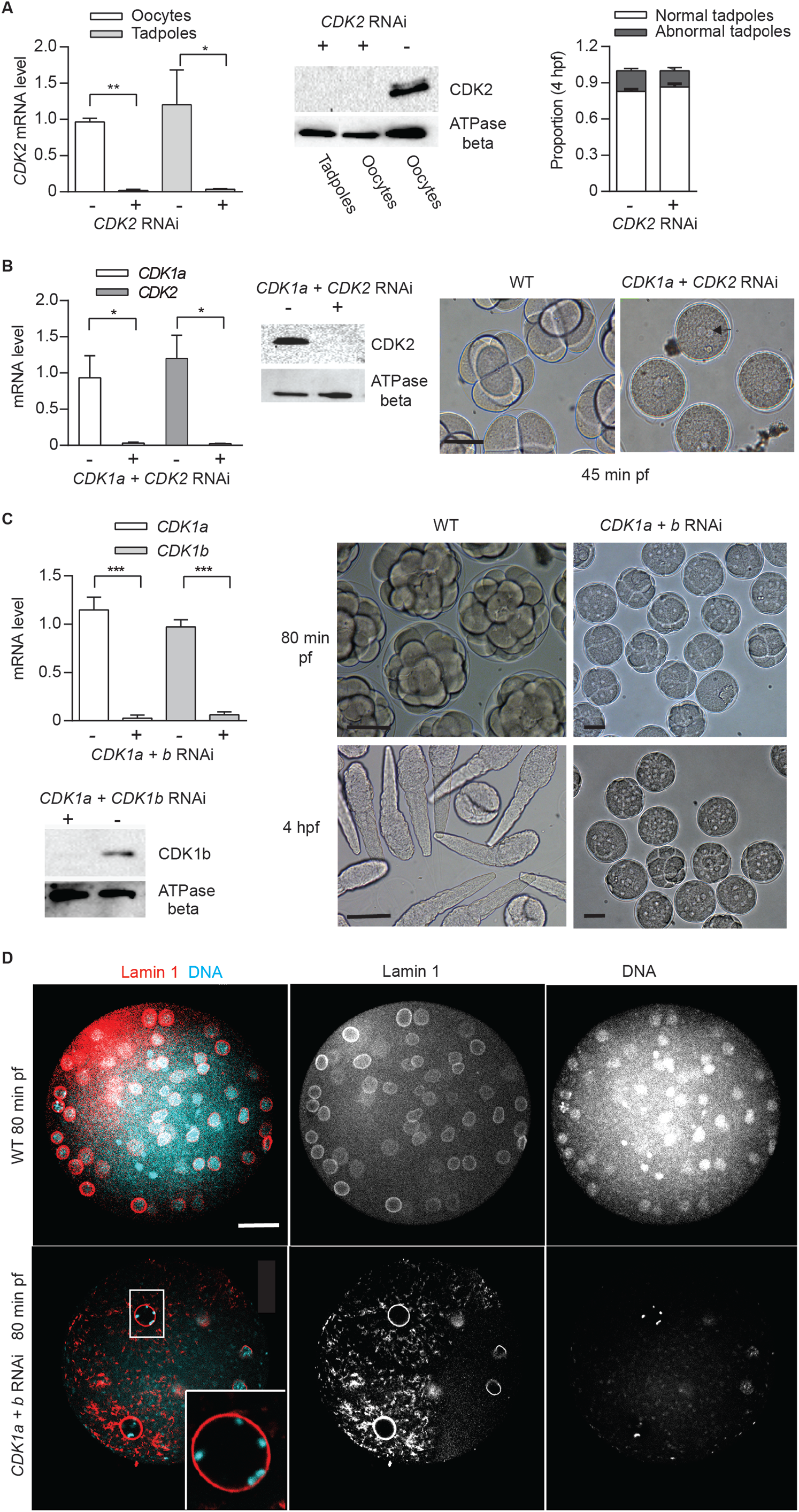
CDK2 is dispensable for *Oikoplerua dioica* embryogenesis. Synergistic defects of *CDK2 + CDK1a* RNAi and *CDK2* + *CDK1b* RNAi in embryonic divisions. (*A*) CDK2 is dispensable for embryonic divisions. Right panel: efficient knockdown of *CDK2* in oocytes and tadpoles, spawned from ovaries injected with dsRNA against *odCDK2* at D4. Mid panel: Western blot showing absence of CDK2 protein in *CDK2* RNAi oocytes and tadpoles. Right panel: no significant differences in developmental outcome between *CDK2* deficient and wild-type embryos. (*B*) Synergistic effect of *CDK2 + CDK1a* RNAi on embryonic divisions. Left Panel: efficient knockdown of *CDK1a* and *CDK2* in oocytes from ovaries that had been injected with dsRNA against *odCDK2* and *CDK1a* at D5. Mid panel: Western blot showing absence of CDK2 protein in CDK1a + *CDK2* RNAi in oocytes. Right panel: at 45 min pf, *CDK1a + CDK2* deficient zygotes were arrested prior to pronuclear fusion. Scale bars: 50 μm. (*C*) Synergistic effect of *CDK1a* + *b* RNAi on embryonic division. Upper left panel: efficient knockdown of *CDK1a* and in oocytes from ovaries that had been injected with dsRNA against *CDK1a* + *b* at D5. Lower left: Western blot showing absence of CDK1b protein in *CDK1a* + *b* RNAi oocytes. Right panel: *CDK1a + b* deficient zygotes finished 3 rounds of slowed divisions before arresting at the 4^th^-5^th^ embryonic division. Scale bars: 50 μm. Where indicated, ***P<0.001, **P<0.01, *P<0.05. (*D*) Confocal images of Lamin 1 and DNA staining of wild-type versus *CDK1a* + *b* RNAi embryos at 80 min post-fertilization. *CDK1a* + *b* deficient zygotes arrest at prophase with super-condensed chromosomes at the nuclear periphery (inset) after the third division. Scale bars: 20 μm.

### Coevolution of *Oikopleura dioica* CDK1d and Cyclin Ba

Vertebrate Cyclin B1 is required for meiosis and mitosis. Most invertebrates, except *C. elegans* (van der Voet et al. 2009), have a single ortholog of vertebrate Cyclin B1/B2 (fig. 1B), and this ortholog is required for meiosis and mitosis (Okano-Uchida et al. 1998). Oscillation of Cyclin B levels orchestrate CDK1 kinase activity during M-phase. As for other invertebrates, all 4 hermaphroditic *Oikopleura* species have a single Cyclin B (Cyclin Bc), whereas dioecious *O. dioica* has Cyclins Ba and Bb (almost identical to Ba) in addition to Cyclin Bc. The *cycBa* gene is located on the X chromosome, and is specifically expressed in ovaries. Knockdown of Cyclin Bc, showed no significant differences in Cyclin Ba levels, H1 kinase activity in oocytes, or embryonic development compared to wild type (fig. 5A), indicating that Cyclin Bc is dispensable for meiosis and embryonic divisions. In contrast, *cycBa* RNAi oocytes were defective for embryonic division (fig. 5B). Therefore, in *O. dioica* oogenic meiosis and early embryonic cleavage cycles, Cyclin Ba was required, whereas Cyclin Bc was dispensable.

**FIG. 5.**
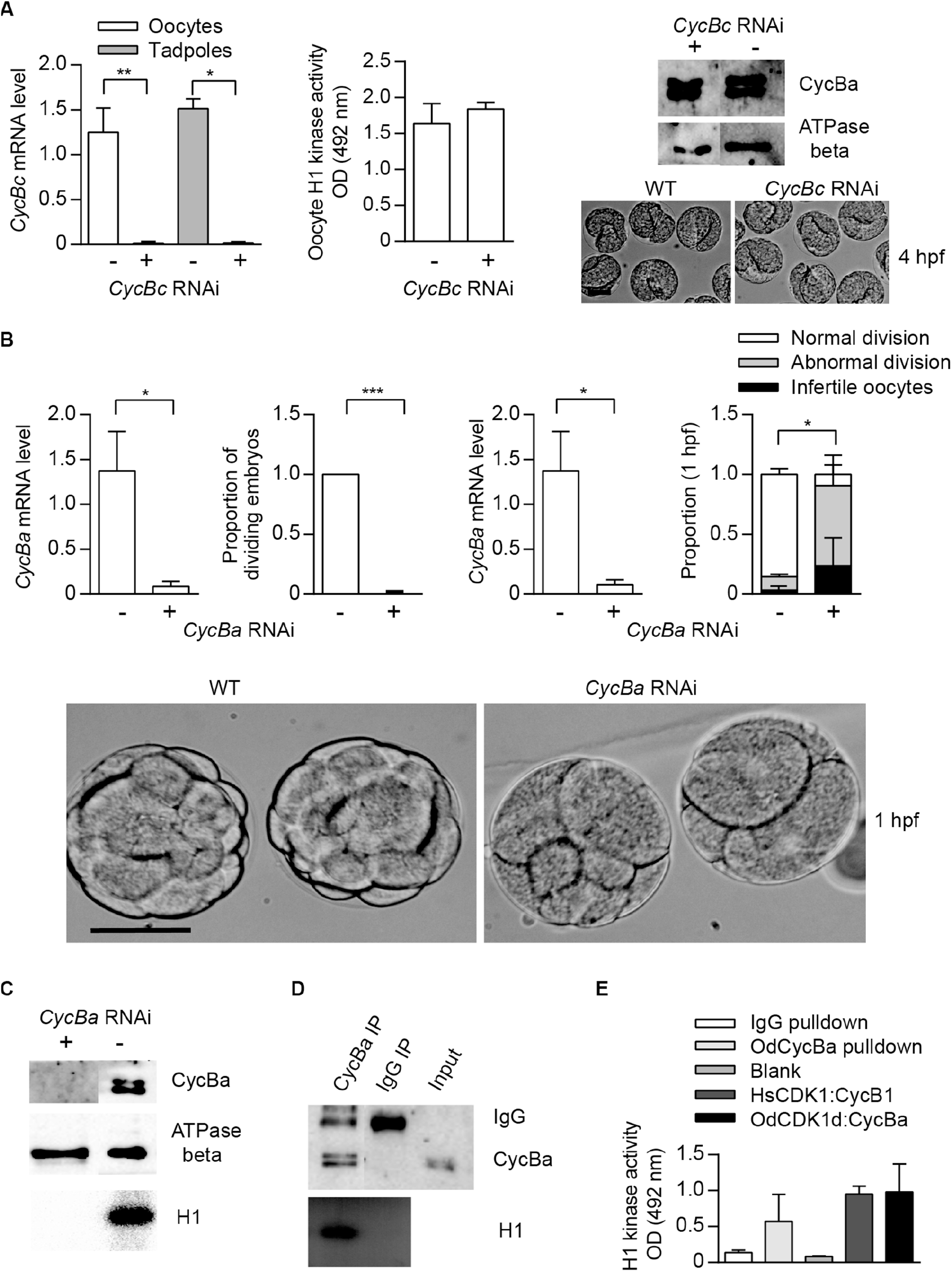
Cyclin Ba, is required for *O. dioica* embryonic cell cycles, whereas Cyclin Bc is dispensable. (*A*) Cyclin Bc is dispensable for embryonic divisions. Left panel: efficient knockdown *cycBc* in oocytes and tadpoles spawned from ovaries that had been injected with dsRNA against *cycBc* at D5. Mid panel: no significant difference in mean H1 kinase activity between Cyclin Bc deficient and wild-type oocytes (metaphase I arrested). Upper right panel: Western blot showing that knockdown of *cycBc* had no effect on protein levels of Cyclin Ba. Lower right panel: images show Cyclin Bc deficient and wild-type embryos had identical phenotypes at the tailbud stage, 4 hpf. (*B*) Efficient knockdown of *cycBa* in oocytes spawned from ovaries that had been injected with dsRNA against *cycBa* at D5. Cyclin Ba deficient oocytes exposed to wild-type sperm generated two embryonic phenotypes: infertile and abnormal divisions. Abnormal division phenotypes, compared to wild-type, are illustrated in the bottom panels at 1 hpf. Where indicated, ***P<0.001, **P<0.01, *P< 0.05. (*C-E*) CycBa-CDK1d complexes have histone H1 kinase activity. (*C*) H1 kinase activity assay was present in wild type oocytes but absent in *cycBa* RNAi knockdown oocytes (Feng and Thompson 20118). (*D*) Cyclin Ba immunoprecipitates from oocytes had H1 kinase activity whereas IgG oocyte immunoprecipitates did not. (*E*) H1 kinase activity levels of *O. dioica* Cyclin Ba pulldowns compared to IgG pulldowns and *in vitro* expressed complexes of Human (Hs) CDK1:Cyclin B1 and *O. dioica* (Od) CDK1d:Cyclin Ba. Scale bars: 50 μm.

Among the oikopleurids, CDK1d and Cyclin Ba specifically appear in *O. dioica* and share several features: X chromosome location, ovary-specific expression, similar degradation dynamics after fertilization (fig. 6B) and similar RNAi phenotypes in meiosis and mitosis. Therefore, we wished to determine if they form an active kinase complex. Cyclin Ba immunoprecipitates possessed H1 kinase activity, whereas lysates from infertile *cycBa* RNAi oocytes lose H1 kinase activity (fig. 5C). This led us to test whether *in intro* constituted odCDK1d-Cyclin Ba complexes possesses kinase activity. GST-odCDK1d recombinants expressed in insect sf9 cells were insoluble. Co-expression with his-Cyclin Ba improved GST-odCDK1d solubility. Purified GST-odCDK1d: his-Cyclin Ba complexes possessed H1 kinase activity (fig. 5E). Surprisingly, the conserved Cyclin B Y170 residue which is critical for CDK1 binding is divergent in odCyclin Ba (replaced by His), whereas it is conserved in odCyclin Bc (supplementary fig. S7). A Y170A substitution is known to completely abolish CDK1:Cyclin B interaction in mouse, human, *Xenopus laevis* and *Ciona intestinalis* (Goda et al. 2001; Bentley et al. 2007; Levasseur et al. 2013; Levasseur et al. 2019). The CDK1d:Cyclin Ba complex also has modified residues at the position of the salt bridge (Goda et al. 2001) (supplementary fig. S7), which is in spatial proximity to Y170. This suggests that CDK1d and Cyclin Ba may have coevolved an exclusive interaction that does not interfere with the more canonical CDK1a/b/c:CycBc interactions.

**FIG. 6.**
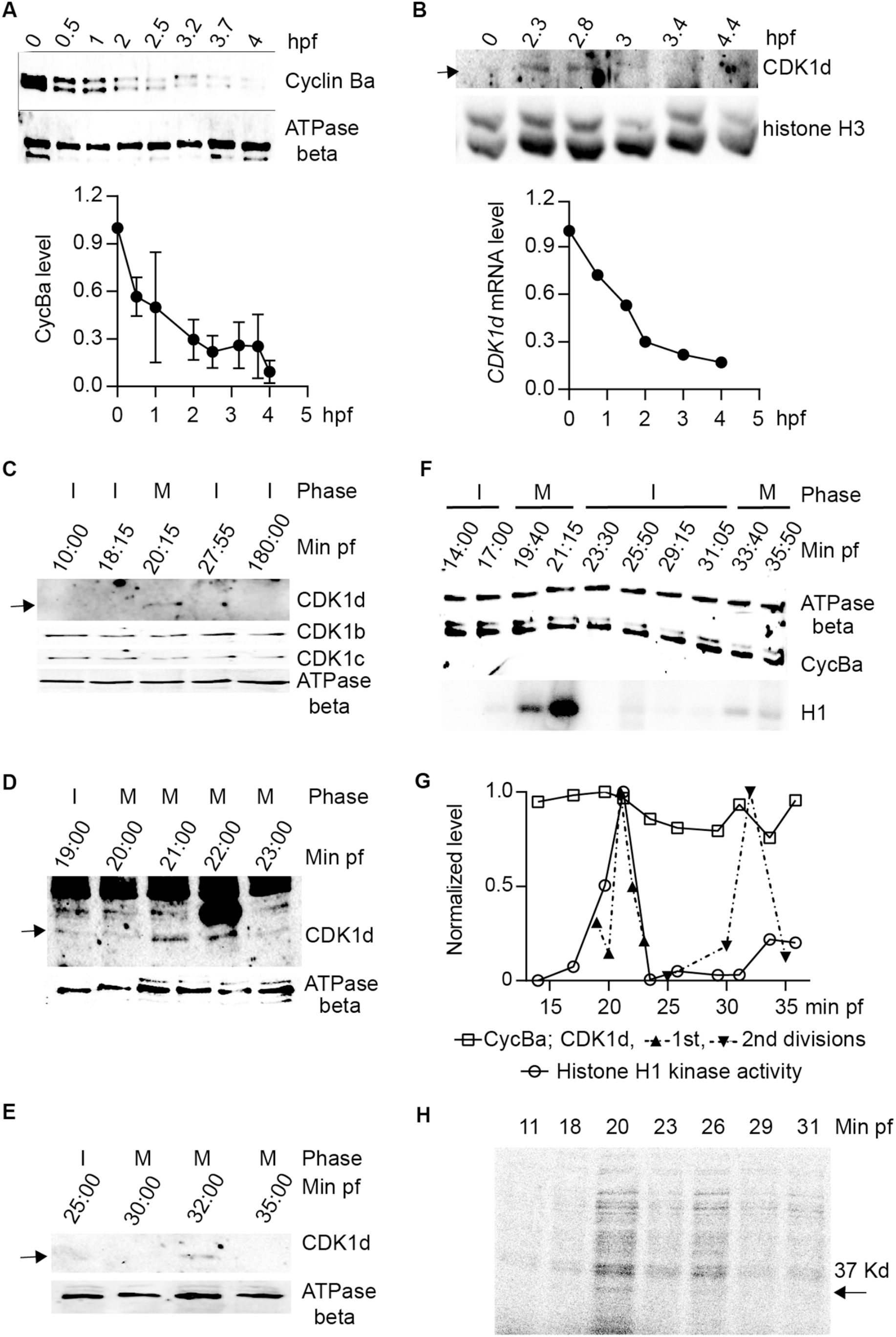
CDK1d levels oscillate during early embryonic cell cycles. (*A*) Cyclin Ba levels do not oscillate but instead, decline steadily during early embryogenesis. Upper panel: Western blot of Cyclin Ba during the first 4 h of development with ATPase beta as a loading control. Lower panel: quantification of Cyclin Ba levels, normalized to levels in unfertilized oocytes. (*B*) Upper panel: Western blot of CDK1d levels during the first 4 h of development with histone H3 as a loading control. Lower panel: *CDK1d* mRNA levels were quantified during early embryonic development by qRT-PCR, normalized to levels in unfertilized oocytes. (*C-E*) CDK1d levels peaked in M phase of first two mitotic embryonic cell cycles. (*C*) Western blot of CDK1b, c, and d levels during the first 3 hpf. Arrow indicates the CDK1d band. I, interphase; M, M-phase. (*D*) Sampling of CDK1d levels at 1-minute internals around M phase of the first embryonic cell cycle. Arrow indicates the oscillation of the CDK1d band. (*E*) Sampling of CDK1d levels around M-phase of the second division cycle. Arrow indicates the oscillation of the CDK1d band. (*F*) Western blot analyses of Cyclin Ba levels and histone H1 kinase activity during the first two embryonic cell cycles. Cyclin Ba levels remained elevated, whereas H1 kinase activity oscillated, with peaks in the first two M-phases. (*G*) Quantification of Cyclin Ba, CDK1d and H1 kinase activity during the first two embryonic cell cycles. (*H*) Oocytes were incubated in artificial sea water with 35S methionine for 1 h before fertilization. Embryos were collected at 3 min intervals during the course of first embryonic cell cycle. The autoradiograph shows *de novo* synthesized protein during the course of this cycle. Arrow indicates the CDK1d band.

### Oscillation of CDK1 levels, instead of Cyclin B levels, orchestrates early embryonic divisions

In marine invertebrates and in *Xenopus* embryos, mitosis is driven by accumulation of Cyclin B to a threshold required to activate CDK1, and mitotic degradation of Cyclin B inactivates CDK1, allowing mitotic exit. Cyclin Ba was stocked in *O. dioica* oocytes and was progressively degraded during early embryogenesis, dropping to low levels by 2.5 hpf (fig. 6A). CDK1d also reached low levels during this same period (fig. 6B). We then performed higher resolution sampling of Cyclin Ba levels at 2 min intervals during the first embryonic cell cycle. Morphologies of the zygotes were recorded at each time point and samples were divided into 2 aliquots and assayed for Cyclin Ba protein levels and H1 kinase activity. Consistent with cell cycle progression, H1 kinase activity oscillated, peaking 20 min post-fertilization, when the nuclear envelope had disappeared and spindles were visible (fig. 6F). H1 kinase levels dropped at 23 min post fertilization, when cleavage furrows formed and dropped to a basal level at 28 min post fertilization, when abscission occurred, and two nuclei were visible. Surprisingly, CDK1d levels oscillated during the first cell cycle (fig. 6 C-G), in contrast to CDK1b and c levels which remained constant. CDK1d levels and H1 kinase activity peaked together at 20 min post fertilization, when NEBD occurred and the spindle assembled. CDK1d levels then peaked again, coincident with a rise in H1 kinase activity during the second cell cycle (fig. 6F and G). To further support this unusual oscillation in CDK1d levels, we labeled oocytes with ^35^S-methionine, and sampled zygotes after *in vitro* fertilization. Autoradiography of SDS-PAGE detected a 37 KD band at the expected CDK1d MW, which appeared 20 min post-fertilization, corresponding to M phase (fig. 6H). Contrary to the oscillation of CDK1d levels, Cyclin Ba levels did not oscillate during the first 2 embryonic cycles (fig. 6H and G), instead, dropping at later embryonic stages. This latter observation is similar to *Drosophila* preblastoderm cycles, where neither bulk levels of Cyclin B, nor the bulk activity of Cdk1 kinase oscillate, with progressive depletion of maternally stocked Cyclin B only leading to oscillations in Cyclin B levels at later developmental stages (Edgar et al. 1994). A key, and novel difference in *O. dioica* is that CDK1d levels did oscillate during these first embryonic cycles.

## Discussion

*Oikopleura dioica* exhibits high intrinsic rates of natural population increase (*r*) (0.68 d–1 to 1.07 d– 1), well above the normal range for a metazoan of its size (Troedsson et al. 2002). The parameter, *r*, is increased by decreasing generation time, and/or by increasing egg number. In *O. dioica*, generation time is highly heritable and genetically correlated with *r*, whereas fecundity is much less heritable and not genetically correlated with *r* (Lobon et al. 2011). This emphasizes the importance of generation time in setting *O. dioica* fitness and is evidence of a life-history trait that has been strongly constrained by evolution (Deibel and Lowen 2012). The short duration of embryogenesis can be a direct target for selection to increase *r*.

We propose a working model for *O. dioica* cell cycle regulation under the framework of multiple CDK1 paralogs during oogenesis and rapid, early embryogenesis (fig. 7). Individual, or combined, knockdowns of CDK1c and e, did not generate any detectable phenotypes. Of note, neither of these CDK1 paralogs was able to complement yeast *cdc28* mutants. Similar to observations in mouse embryos and cultured vertebrate cells, CDK2, the only CDK with a canonical PTSAIRE helix in *Oikopleura*, was dispensable for completion of meiosis and early embryonic divisions, as *CDK2* RNAi zygotes divided and hatched normally (fig. 4A). On the other hand, normal early embryonic divisions required 3 CDK1 paralogs (figs. 4, 5, 7 and supplementary fig. S5), none of which contain a canonical PSTAIRE helix (fig. 1A). Depletion of CDK1a or b alone, delayed embryonic divisions (fig. 3A and B), whereas depletion of CDK1d abolished meiotic NEBD (Ovrebo et al. 2015), disrupted meiosis completion and caused abnormal embryonic cell cycles (fig. 3C-G). Knockdown of CDK1d also depleted oocytes of histone H1kinase activity (Feng and Thompson 2018), as did knockdown of its binding partner, Cyclin Ba (fig. 5C). Double knockdowns of CDK2 and CDK1a arrested zygotes at the pronuclear stage (fig. 4B) whereas double knockdowns of CDK1a and b generated multinucleate embryos with irregular cell division patterns that failed to hatch (fig. 4C and D). CDK1d had critical functions in completion of meiotic M-phases and early embryonic divisions, during which it showed an unusual oscillation of protein levels that peaked in M-phase and decreased during interphase. CDK1d protein dynamics correlated well with the peak of H1 kinase activity and spindle formation during M phase (fig. 6C-H).

**FIG. 7.**
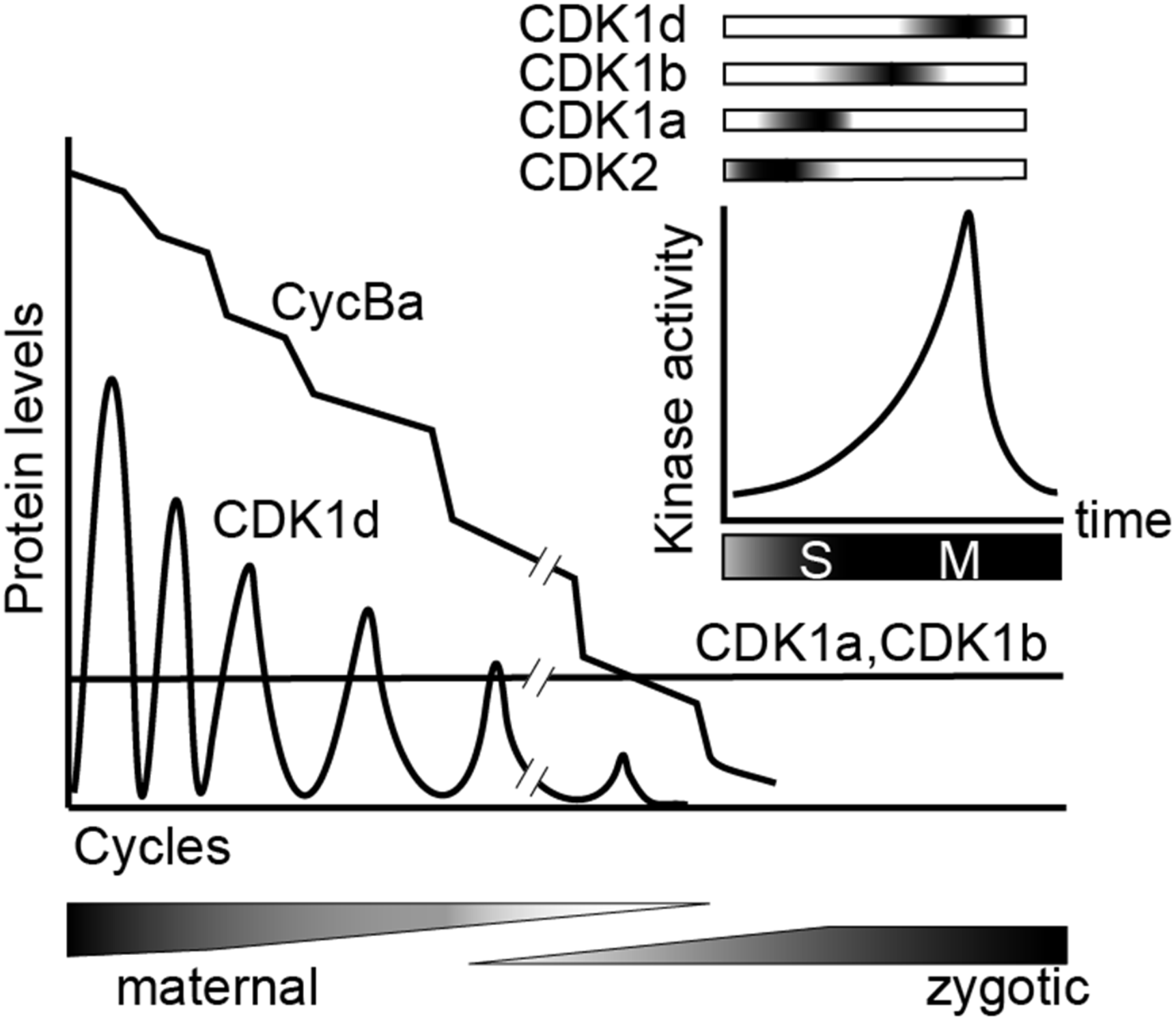
Sequential activation of CDK1a, b and d, and switching from oscillations of CDK1d levels to oscillation of Cyclin Ba levels, drive rapid early embryonic cell cycles in *O. dioica*. Knockdowns revealed that sequential activation of 3 *O. dioica* CDK1 paralogs, a, b and d, are required for proper execution of embryonic divisions leading to embryo hatching. Cyclin Ba, the partner of CDK1d, is maternally stocked and gradually degrades over the first embryonic cycles, whereas unusually, it is CDK1d protein levels that initially oscillate to regulate rapid M-phases. CDK1a and b levels remain constant during this period. During the maternal to zygotic transition, the cell cycle slows and divisions become less synchronous. In the coenocystic ovary, polyploid nurse nuclei are afforded the time to produce maternally stocked transcripts, many arising from genes containing multiple introns of variable length, with the genes often organized in polycistronic operons (Danks et al. 2015). During the period of rapid embryonic cell divisions, many zygotically activated genes are monocistronic, and contain short introns, or are intronless.

The 3 CDK1 paralogs are sequentially activated in a tiling pattern when embryos traverse late S- to M-phase, suggesting a further subdivision of CDK1 tasks and substrate targeting, beyond the CDK1-CDK2 subdivision that occurred much earlier in metazoan evolution. An additional, interesting feature of the *O. dioica* CDK1 paralog complement, is that it is CDK1d protein levels rather than its partner, Cyclin Ba, the only essential Cyclin B for *O. dioica* oogenic meiosis and early embryonic cell cycles, which oscillates during early embryonic cell cycles (fig. 6F and G). This contrasts the generally universal oscillation of Cyclin B levels on a background of stable CDK1 protein levels characteristic of metazoan cell cycle regulation.

During the maternal to zygotic transition, initial, rapid cell cycles, followed by progressive slowing, are a widespread occurrence in a number of species, indicating a consistent benefit to this strategy throughout the evolution of animals with externally deposited eggs (Farrell and O’Farrel 2014; Yuan et al. 2016). This strategy converges on modulating the activity of the cell cycle driver, CDK1: weak inhibitory phosphorylation of CDK1 drives rapid cycles without gap phases, and when inhibitory phosphorylation is introduced, catalyzed by Wee and Myt1 kinases, the cell cycle slows and gap phases are inserted (Tsai et al. 2014). *O. dioica* and, the model arthropod, *Drosophila*, are characterized by a number of similarities with respect to oogenesis and early embryogenesis. Both utilize a syncytial strategy during oogenesis (albeit with significant differences in details), where polyploid nurse nuclei generate transcripts that code for materials stocked in the oocytes, which house largely transcriptionally quiescent meiotic nuclei. Both generate excess Cyclin B protein storage in oocytes, with progressive Cyclin B destruction during succeeding embryonic cycles. They diverge, however, in adaptations of rapid mitotic cycles to Cyclin B preloading. *Drosophila* syncytial embryos adapt Cyclin B preloading to mitosis without cytokinesis, compatible with little oscillation of CDK1 kinase levels due to localized Cyclin B destruction (Edgar et al. 1994). In contrast, completion of cytokinesis during the rapid mitoses of *O. dioica* embryonic cycles appears to require oscillation of CDK1 kinase levels (fig. 6F and G). CDK1d oscillations may be an adaptation to the coenocystic ovary structure and Cyclin Ba preloading. Another possible benefit to storage of Cyclin Ba, rather than CDK1d, could be that progressive destruction of Cyclin Ba by the anaphase promoting complex during embryogenesis would deplete Cyclin Ba, facilitating switching of key cell cycle regulators after hatching. In this regard, regulators, such as CDC25, Wee1, and Emi2, critical in controlling important embryonic cell cycle transitions in other organisms (Farrel and O’Farrell 2014; Tsai et al. 2014; Yuan et al. 2016), are all duplicated in the *O. dioica* genome.

Coenocystic oogenesis is a common feature of the Oikopleuridae (Ganot et al. 2006), allowing synchronous oocyte growth and maturation as an adaption to a semelparous, opportunistic life style. The oikopleurid lineage also shares the presence of CDK1a, b and c paralogs. This raises questions as to the relationship between the specific gain of the CDK1d-Cyclin Ba complex in *O. dioica*, their linkage on the recently evolved X chromosome, and their ovary-specific expression. This pattern could be explained by resolution of sexual antagonism by gene duplication (Connallon and Clark 2011) and higher linkage disequilibrium on the X chromosome (Schaffner 2004). For genes that function both in hermaphrodite ovary and testes, the coenocystic ovary structure evokes conflict between different transcriptional regulation of those genes in polyploid nurse nuclei in the ovary, versus diploid germline progenitor cells in the testis. CDK1c does not show sexually biased gene expression (fig. 2A) (Campsteijin et al. 2012). CDK1c duplication giving rise to CDK1d and e in *O. dioica* may have mitigated sexual antagonistic conflict by allowing CDK1d (sexually biased expression to ovaries) to evolve essential ovarian functions. The X chromosome location of CDK1d might facilitate CDK1d ovary-specific expression since duplicates beneficial to females preferentially accumulate on the X chromosome where they are exposed to selection twice as often in females versus males (Connallon and Clark 2011). Indeed, when compared to zygotic promoters, maternal promoters in *O. dioica* are located on the X-chromosome more frequently than expected, (χ2 = 43.34, df = 1, p = 4.61 × 10^−11^), revealing a female-bias of X-linked genes in *O. dioica* (Danks et al. 2018). X-linkage of *CDK1d* and *Cyclin Ba* loci would then efficiently sustain linkage disequilibrium, reinforced through selection on the recently formed X chromosome (Bachtrog et al. 2009). Since there will be no recombination in the heterogametic male, this would contribute to an increase in sweeping to fixation in the population.

In *O. dioica*, the 3 CDK1 paralogs (a, b and d), which are required for correction execution of early embryonic cell cycles, exhibit sequence diversification on their activation segments, ATP binding sites and Cyclin B interaction interfaces. This raises the possibility of different k_cat_/k_m_ catalytic outputs or phosphorylation site specificities during these rapid cell cycles. With respect to the K89 residue in the hinge region, both CDK1a and b, which participate in events preceding CDK1d, share a K89E/D substitution. This substitution would likely reduce CDK1 k_cat_. CDK1a, b and d also have non-overlapping activation segments, suggesting CDK1 paralogs may have different affinities toward the same substrate phosphorylation site and/or different phosphorylation site preferences. The sequential activation of CDK1a, b and d may provide a more stepwise, as opposed to progressive, increase of kinase activity, which may enhance and sharpen transitions towards early, middle and late substrates across early embryonic cell cycles. In metazoan cell cycles, the division of labor between CDK1 and CDK2 is not absolute. CDK1 retains essential roles in late S phase that CDK2 does not efficiently complement (Katsuno et al. 2009; Nakanishi et al. 2010; Farrell et al. 2012). Inactivation of Cdk1 in temperature-sensitive Cdk1 mutant cell lines (FT210) resulted in a prolonged S phase accompanied by ineffective firing of late replicon clusters and reduced the density of active origins during late S phase (Katsuno et al. 2009). In *Drosophila* pre-mild blastula transition S phases, it is Cdk1 activity, rather than Cdk2 activity that regulates late replication (Farrell et al. 2012; Seller and O’Farrell 2018). The requirement for CDK1 in late S phase, coexists with its need to be repressed during early S phase (Larochelle et al, 2007) and its control in late S phase by Chk1 (Lemmens et al. 2018), to prevent premature triggering of M phase (Szmyd et al. 2019). In *O. dioica*, the conflict of CDK1’s roles in the metazoan cell cycle of regulating both late S and M phase, might be relaxed by CDK1 duplication and subsequent partitioning of ancestral CDK1’s roles in S and M phase to specific paralogs. Our results indicate that CDK1a collaborates with CDK2 to drive S phase and suggest that this paralog may exhibit reduced risk of triggering M phase, through lower intrinsic kinase activity, arising from substitution in hinge region (supplementary fig. S2). On the other hand, the CDK1d paralog has become a specialist of M phase events without additional constraints imposed by regulation of late DNA replication. This may reflect adaptation to very rapid embryonic cell divisions in this planktonic organism.

A conformation change of the very highly conserved PSTAIRE helix is central to CDK1 inactivation/activation transitions (Huang et al. 2012). It is therefore surprising that this sequence was modified in all *O. dioica* CDK1 paralogs. *O. dioica* CDK2, the only CDK in this species with a conserved PSTAIRE, was dispensable for embryonic divisions, recalling the situation in plants, where the CDK1 ortholog, CDKA, which has the canonical PSTAIRE, is dispensable for the cell cycle, whereas CDKB, deviating in the PSTAIRE, has essential functions in the absence of CDKA (Nowack et al. 2012). Strikingly, the modified PSTAIRE of the *O. dioica* CDK1d paralog (PPTSLRE) shows convergence over great evolutionary distance with plant CDKB sequences (P[P/S]T[A/T]LRE), and in both *O. dioica*, and plants, these variants exhibit increased specialization to M-phase (Joubes et al. 2000; Nowack et al. 2012). The suite of naturally evolved *O. dioica*, CDK1:Cyclin B complexes, with critical residue substitutions in their activation segments, ATP binding sites, CDK1-Cyclin B interaction interfaces, and Cyclin B-substrate interaction elements, provide a fertile foundation for structural studies of how these changes elicit enhanced kinase selection for subsets of G2-M, CDK1 cell cycle substrates.

## Materials and Methods

### Animal culture and collection

*O. dioica* were maintained in culture at 15°C (Bouquet et al. 2009). Day 4–6 animals were placed in filtered seawater, removed from their houses and anesthetized in cold ethyl 3-aminobenzoate methanesulfonate salt (MS-222, 0.125 mg/ml; Sigma) before collection.

### *In vitro* fertilization (IVF) and collection of synchronously dividing zygotes

Sperm solutions were prepared by transferring 5 mature males to a petri dish containing 10 ml artificial seawater, on ice, and sperm quality was assessed using a Nikon Eclipse E400 microscope. Mature females were transferred to artificial sea water in 24-well plates coated with 0.9% agarose, artificial sea water was changed every hour. Oocytes were fertilized with 400 μl sperm solution and fertilized oocytes were washed in 8 ml artificial sea water within 50 seconds. Synchronicity of fertilization was verified by following polar body exclusion and early cleavages.

### Phylogenetic Analyses

A search of larvacean CDK1, CDK2 and Cyclin B proteins was performed using TBLASTN against genomes and transcriptomes of *Oikopleura albicans, O. longicauda, O. vanhoeffeni and Fritillaria borealis*. *O. dioica* CDK1, CDK2 and Cyclin Bc were used as queries with E-value cutoff set to 1e-005. *Ciona intestinalis* and *Branchiostoma floridae* proteins were obtained from the JGI Genome Portal (https://genome.jgi.doe.gov/mycocosm/home). For identified targets, amino acid sequences of the N-terminal cyclin box or the kinase domain of CDK1/2 were aligned (clustalX). Alignments were manually edited (Bio-edit), and secondary structure analysis (psipred) was used to guide alignments. Phylogenetic relationships were inferred using maximum likelihood (ML) using MEGA7 (Kumar et al. 2016), with amino acid sequence alignment based on Jones-Taylor-Thornton (Jones et al 1992). Estimates of the gamma-shape parameter and proportion of invariant sites were then used to obtain the 500 bootstrap replicates.

### Yeast Complementation

We used a W303a-background yeast wild-type strain SCU893 (*MATa ura3 ade2-1 trp1-1 leu2-3112 his3-11 bar1::hisG*) and its derivative SCU112 (*cdc28-4*). To complement a *cdc28* mutation of *S. cerevisiae*, the open reading frames of the *O. dioica CDK1* paralogs and *CDK2* cDNAs were cloned in pCR™4-TOPO® TA Vector (Thermofisher). cDNA fragments were isolated by digestion with *EcoR*I and *Not*I and ligated into the yeast pYES2 expression vector (Invitrogen) downstream of the inducible GAL1 promoter. The resulting CDK1 paralog plasmids were individually transformed by the lithium acetate method (Ito et al. 1984) into the *S. cerevisiae cdc28-4* strain. As a control, *cdc28-4* was transformed with the empty pYES2 vector. The transformants were grown for 4 days at 25°C, and the resulting colonies were cultivated overnight at 25°C in CM-ura medium containing glucose as a carbon source. The cells were pelleted, washed, resuspended in CM-ura medium containing 2% glycerol, and grown further at 25°C. After 4 h, the cells were pelleted and resuspended in CM-ura medium containing either glucose (2%), or galactose (2%)–raffinose (1%) as carbon source, and grown at 25°C, or the restrictive 36°C, respectively.

### Antibodies

Rabbit polyclonal affinity–purified anti-Cyclin Ba/b (acetyl-NRDLNIQESGPVKAVVNAC-amide; acetyl-CLEFLRRFSRVAEETIDPKEY-amide), anti-CDK1b (acety1-CPDFTKPSTLNVNTSLNEMDM-amide), anti-CDK1c (acetyl-MRKRNIDRPPTQDLNNYC– amide), ant-Cdk1d (acety1-MIKAKSGTQDLSNYKRC-amide), and anti-lamin1 (acetyl-QSPISLPPLSGSTC-amide) were from 21st Century Biochemicals (Marlboro, MA). Other antibodies: anti-PSTAIR (Abcam, ab10345), anti-ATPB (Abcam, ab37922), anti-histone 3 (Abcam, ab10799). Secondary antibodies for immunofluorescence: goat anti-rabbit IgG (H+L) Alexa Fluor 568 (Thermofisher, A-11011); and for western blots: goat anti-rabbit IgG (H+L) superclonal™ secondary antibody, HRP (Thermofisher, A27033), Peroxidase-AffiniPure Goat Anti-Mouse IgG (H+L) (Jackson ImmunoResearch, 115-035-003).

### Quantitative Reverse Transcription–PCR, Immunofluorescence and Western Blots

Total RNA was isolated from snap-frozen animals. For qRT-PCR, 1 μg of purified total RNA was subjected to RT using M-MLV reverse transcriptase (Invitrogen). The quantitative polymerase chain reaction (qPCR) (CFX-96; Bio-Rad) was performed using cDNA templates synthesized from an equivalent of 5 ng total RNA, 10 μl of qPCR 2 Master Mix (Bio Rad), and 500 nM primers in a total volume of 20 μl. After initial denaturation for 5 min at 95 °C, 45 cycles of 95 °C for 15 s, 60 °C for 20 s, and 72 °C for 20 s were conducted, with a final extension for 5 min at 72 °C. RT-negative controls were run to 45 cycles. RPL23 and EF-1β mRNA levels served as normalization controls in all qRT-PCRs. For RNAi experiments, target mRNA levels were normalized to control zygotes. Primer pair specificities were confirmed by melting curve evaluation, cloning, and sequencing of amplicons. In these experiments, mean values normalized to EF-1β control transcripts are presented. Immunofluorescence and Western blots were performed as previously (Campsteijn et al. 2012).

### Microinjection

Gonad microinjections were performed as previously (Ovrebo et al. 2015). Injection solutions were prepared by mixing 300 ng/μl double stranded RNA (dsRNA) in RNase free PBS. Anesthetized day 4 animals in filtered seawater were transferred to Petri dishes coated with 2% agarose. Gonads were injected on a Nikon Eclipse TE2000-S inverted microscope equipped with Narishige micromanipulators and transjector 5246 micro injector (Eppendorf). Injected day 4/5 animals were transferred to watch glasses containing filtered seawater for recovery and then transferred to 6 L beakers for subsequent culture. Animals were cultured until spawning for in vitro fertilization (IVF) and immunofluorescence. Before oocyte injection, oocytes were transferred to 24-well plate coated with 0.9% agarose containing fresh artificial seawater (Red Sea) for 1 h. Samples of 50 oocytes were used for qRT-PCR. Remaining oocytes were fertilized and developmental defects arising from various treatments were assessed by unpaired Student’s t-test for equal variances, or, unpaired, Welch’s t-test for unequal variances. Effect size was calculated as Cohen’s *d* = 2*t*/SQRT (*df*).

### *In vitro* CDK1 kinase activities

CDK1 kinase activities were determined as previously (Feng and Thompson 2018). Equal numbers of oocytes were snap frozen and thawed twice. Cyclin Ba immunoprecipitates, purified his-Cyclin Ba-GST OdCdk1d complexes were aliquoted, kept at 80°C and thawed before use. Samples were mixed with biotinylated MV peptide and 1 mM ATP incubated at 30°C for 30 min. After stopping the phosphorylation reaction, ELISA was performed using anti-phospho MV peptide antibody (MESACUP) and the CDK1 Kinase Assay Kit (MBL). Finally, POD-conjugated streptavidin was used to detect phosphorylated MV peptide and color development measured at 492 nm. Differences in CDK1 kinase activities from different knockdown treatments were assessed by the Student’s T-test. H1 kinase assays were performed as previously (Levasseur and McDougall 2000). After thawing snap frozen oocyte samples on ice, 2 μl of 6× kinase reaction mixture (pH 7.2) containing histone H1 (Sigma type III from calf thymus), 0.6 mM ATP, 0.5 mCi/ml ^[32P]^ATP, and 60 μM cAMP-dependent protein kinase inhibitor was added to 10 μl of oocyte sample. The reaction was started by transfer to a 30°C heat block for 30 min and stopped by adding 5× Laemmli buffer. The samples were then heated to 80°C for 10 minutes and resolved on 15% polyacrylamide gels. Gels were placed in a Phosphorimager (Fujix; 1500 Bas Reader) and incorporation of [^32^P] was quantified.

### Metabolic labeling of zygotes

Unfertilized oocytes obtained from individual females were transferred into one well of 24-well plate coated with 0.9% agarose. After addition of a mixture of ^35^S-labeled amino acids (Pro-mix, Amersham) to a final concentration of 330 μCi/ml and incubation for 60 min at room temperature, oocytes were fertilized and harvested at different times post fertilization. The samples were briefly centrifuged to eliminate excess of labeled mixture and 5× Laemmli buffer was added to the pellets. Samples were then heated to 80°C for 10 min and resolved on 10% polyacrylamide gels. Gels were stained with 0.5% Coomassie blue R250 in 45% methanol, 45% water, 10% acetic acid, and destained in 20% methanol, 75% water, 5% acetic acid; and dried on Whatman 3MM paper. Dried gels were autoradiographed using a Phosphorimager (Fujix; 1500 Bas Reader).

### Capped mRNA and dsRNA synthesis

For indestructible Cyclin Ba injection, site directed mutagenesis (New England Biolabs) was used to delete the destruction box (NTEREQLQALN) using a cyclin Ba/b entry clone as template with subsequent recombination with pSPE3-Venus-RfA. The resulting plasmid was used as a template for PCR amplification. Amplicons were purified by phenol-chloroform extraction followed by ethanol precipitation. Capped mRNAs were synthesized from these templates using mMessage mMachine (Ambion) followed by poly(A) tailing (Ambion) according to the manufacturer’s protocol and purified by Lithium chloride precipitation. DNA templates for dsRNA synthesis were prepared by PCR using gene-specific primers with T7 overhang and purified by phenol-chloroform extraction. Sense and antisense RNA were synthesized in a single reaction following the recommended protocol using T7 RiboMAX™ Express RNAi system (Promega).

## Acknowledgments and funding information

We thank Haiyang Feng, for helpful discussion and A. Aasjord and K. N. Nøkling for providing animals from the *Oikopleura* culture facility. Yeast strains were provided by Jorrit Enserink, Univ. of Oslo. We thank the Chourrout S1 group at the Sars Centre for early access to sequence data from hermaphroditic appendicularian species. *Oikopleura dioica* genome and transcriptome are available at http://www.genoscope.cns.fr http://oikoarrays.biology.uiowa.edu/Oiko/ and transcription start site CAGE data at the NCBI Gene Expression Omnibus https://www.ncbi.nlm.nih.gov/geo/ under accession number GSE78794. The *Fritillaria borealis* genome is available at GenBank https://www.ncbi.nlm.nih.gov/genbank/: SDII00000000 and other *Oikopleura* species genomes at GenBank SCLD01000000 to SCLH01000000. This work was supported by grants 183690/S10 NFR-FUGE and 133335/V40 from the Norwegian Research Council (E.M.T.).

## Author Contributions

X.M., and E.M.T. conceived and designed the experiments. X.M. performed experiments and X.M. and E.M.T. analyzed the data. J.I.Ø performed CDK1-GFP cmRNA injection and immunostaining. X.M. and E.M.T. wrote the manuscript. All authors approved the manuscript.

