## Supplemental material for "Evolution of CDK1 paralog specializations in a lineage with fast developing planktonic embryos"

### SUPPLEMENTARY INFORMATION

*Evolution of multiple CDK1 paralogs towards specializations in late cell cycle events in a lineage with fast developing planktonic embryos*

Xiaofei Ma[1], Jan Inge Øvrebø[1,2#], Eric M Thompson[1,2 *]

[1]Sars International Centre, and [2]Dept. of Biological Sciences, University of Bergen, Bergen, Norway

[#]Present address: Huntsman Cancer Institute, University of Utah, USA

#### SUPPLEMENTARY METHODS

##### Test of Selection

Branch-sites tests of selection are phylogeny-based tests that require the *a priori* identification of a foreground branch that is hypothesized to be evolving at a different rate than the background branches in the rest of the tree. Codons are alignment with PRANK, gaps in alignment were deleted. We first set a series of nested models specifying particular branches as foreground branches: the lineage subtending the *CDK1a* clade, the lineage subtending the *CDK1b* clade, the lineage subtending the *CDK1c* clade. Using a likelihood ratio test, we then tested the hypothesis, that a model which included one class of sites with $\omega > 1$, indicating positive selection, was superior to a model where $\omega = 1$. We estimated codon frequencies on the basis of observed nucleotide frequencies at each of the three codon positions, as a procedure to reduce false positives due to potential nucleotide biases. We also used the empirical Bayes method as implemented in PAML 4 (Yang 2007) to identify the positively selected sites along the 3 branches. The synonymous (silent) and non-synonymous (non-silent) nucleotide substitution rates of each branch were also estimated by the modified Nei–Gojobori method (Zhang et al. 1998), based on tree topology inferred by maximum likelihood (ML) using MEGA (Kumar et al. 2016).

##### Homology Modeling

Active and inactive hsCDK1 was predicted with I-TASSER (Yang et al. 2015), using the hsCDK1-cyclin B crystal structure (Brown et al. 2015). 3D models were visualized using Pymol (DeLano 2002). Based on alignment of odCDK1 paralogs and hsCDK1, nonconservative variations between odCDK1s and hsCDK1 were colored in the models.

##### Recombinant protein expression in insect cells

Coding regions of Cyclin Ba/b and OdCdk1d were cloned into pCR™8/GW/TOPO and recombined into pDEST™10 (N-terminal $His_6$-TEV fusion tag) or pDEST™20 (N-terminal GST fusion tag), according to manufacturer's (Thermofisher Scientific) instructions. The EMBL Protein Expression and Purification Core Facility performed protein expression and purification. The pDEST10_Cyclin Ba/b and pDEST20_OdCdk1d plasmid DNA were used for transposition into *E. coli* DH10EMBacY cells. Bacmid DNA was isolated from a 3 ml overnight culture of positive (white) colonies according to the MultiBac protocol (Nie et al. 2013). The bacmid DNA was mixed with X-tremeGENE transfection reagent and Sf-900 III SFM medium and the mixture was added to Sf21 insect cells. After 3 days, the $V_0$ virus was harvested and used to infect a larger Sf21 culture for $V_1$ virus generation. For co-expression of $His_6$-tagged Cyclin Ba/b and GST-tagged OdCdk1d, 500 ml Sf21 insect cells was co-infected with 2 ml Cyclin Ba/b $V_1$ virus and 2 ml OdCdk1d $V_1$ virus and cultured for 72 h at 27°C. Cells were harvested by centrifugation and resuspended in running buffer (50 mM Tris pH 8.0, 250 mM NaCl and 20 mM imidazole) supplemented with benzonase and cOmplete EDTA-free protease inhibitors. Cells were lysed using a Dounce homogenizer and the lysate was cleared by centrifugation (10000 x *g*, 30 min,

4°C). The cleared lysate was incubated with 200 µl Ni-NTA (Qiagen) beads for 2 h at 4°C. After incubation, the mixture was transferred to an empty PD10 column, the beads were washed with running buffer and then eluted in 200 µl fractions with running buffer supplemented with 300 mM imidazole. All fractions were analysed by SDS-PAGE and the fractions containing the Cyclin Ba/b–OdCdk1d complex were concentrated, flash-frozen in liquid nitrogen and stored at -80°C until further use.

#### SUPPLEMENTARY TABLES

**Table S1. CODEML analyses of selection among *Oikopleura CDK1* paralogs.**

| Branch site models | Site class | Proportion | ω Background | ω Foreground | Ln L | 2 Δ L[a] | Positively selected sites[b] NEB prob (ω >1) | BEB prob (ω >1) |
|---|---|---|---|---|---|---|---|---|
| | 0 | 0.90440 | 0.01779 | 0.01779 | | | N/A | N/A |
| Model A | 1 | 0.01233 | 1.00000 | 1.00000 | -9374.15 | | | |
| null | 2a | 0.08215 | 0.01779 | 1.00000 | | | | |
| CDK1a | 2b | 0.00112 | 1.00000 | 1.00000 | | | | |
| Model A | 0 | 0.90524 | 0.01850 | 0.01850 | | 2,7649 | 25 T 0.970* | 61 P 0.889 |
| alternative | 1 | 0.01253 | 1.00000 | 1.00000 | | (P = 0.0481)* | 61 P 0.994** | 85 M 0.962* |
| CDK1a | 2a | 0.08111 | 0.01850 | **41.01038** | -9372.77 | | 85 M 0.999** | 200 K 0.836 |
| | 2b | 0.00112 | 1.00000 | **41.01038** | | | 104 L 0.983* | |
| | | | | | | | 178 S 0.981* | |
| | | | | | | | 194 F 0.954* | |
| | | | | | | | 200 K0.991** | |
| | 0 | 0.91289 | 0.01717 | 0.01717 | | | N/A | N/A |
| Model A | 1 | 0.01462 | 1.00000 | 1.00000 | | | | |
| null | 2a | 0.07134 | 0.01717 | 1.00000 | -9803.76 | | | |
| CDK1b | 2b | 0.00114 | 1.00000 | 1.00000 | | | | |
| | | | | | | 7.5220 (P = 0.0030)** | | |
| Model A | 0 | 0.93943 | 0.01735 | 0.01735 | | | 21 G 0.989* | 21 G 0.843 |
| alternative | 1 | 0.01303 | 1.00000 | 1.00000 | | | 68 D 0.977* | 114 Q 0.970* |
| CDK1b | 2a | 0.04689 | 0.01735 | **28.14003** | -9800.00 | | 114 Q 0.998** | 154 G 0.875 |
| | 2b | 0.00065 | 1.00000 | **28.14003** | | | 154 G 0.992** | |
| Model A | 0 | 0.92043 | 0.01727 | 0.01727 | | | N/A | N/A |
| null | 1 | 0.01183 | 1.00000 | 1.00000 | | | | |
| CDK1c | 2a | 0.06688 | 0.01727 | 1.00000 | | | | |
| | 2b | 0.00086 | 1.00000 | 1.00000 | -9377.39 | | | |
| | | | | | | 7,8597 (P = 0.0020)** | | |
| Model A | 0 | 0.92081 | 0.01831 | 0.01831 | | | 105 V 0.996** | 183T 0.858 |
| alternative | 1 | 0.01200 | 1.00000 | 1.00000 | | | 107 S 1.000** | |
| CDK1c | 2a | 0.06632 | 0.01831 | **999.00000** | -9373.46 | | 117 V 0.992** | |
| | 2b | 0.00086 | 1.00000 | **999.00000** | | | 182 S 0.968* | |
| | | | | | | | 183 T 1.000** | |

Positively selected sites in CDK1a and b branches were identified by the empirical Bayes method implemented in PAML 4.1. [a]Branch-site model A is specified using the model 2 NSsites = 2, LRT = 2Δl, df = 1, with significances (*P<0.05; **P<0.01) determined by one-sided Chi-squared tests. [b]Only positively selected sites with naive empirical Bayes (NEB) or Bayes empirical Bayes (BEB) posterior probabilities (prior distribution estimated from the data) equal or greater to 95% (*), or 99% (**), are indicated. Numbering and residues based on human CDK1. N/A, not applicable.

**Table S2. Primer sequences for cyclin, CDK and reference genes**

| Target transcript | Primer | Primer Sequence (5’ to 3’) |
|---|---|---|
| **Cyclins** | | |
| Cyclin Ba/b | CCQ49F | GCAAGTTTACGGCTACTTGCGCTT |
| | CCQ50R | ATCCATGTACGCGACGGTGAGAAA |
| Cyclin Bc | CCQ55F | AAGGGAATCGGTGGTATTGCCAAG |
| | CCQ56R | AGCATTTGAGCTCGGGTGGTTCTA |
| **CDKs** | | |
| CDK1a | CCQ393F | CTCATTTTCGAGTTTCTTTCGTGC |
| | CCQ394R | AGCCCTCCTTCCAGTCCTTG |
| | CCQ395F | GCCGAAATAGCAACGCTCAGA |
| | CCQ396R | TTTCCTAAGTCGCACATTGGC |
| CDK1b | CCQ401F | CGCAAGATCGTCAGAAGCAGA |
| | CCQ402R | CTCGACCAAAATCGTGATCAGATC |
| | CCQ403F | GCAATGGGTATCAAGAAACGTCTG |
| | CCQ404R | GCTGAAACGCCTTCCGCA |
| CDK1c | CCQ409F | ATCGACGACCAGAAAGCCG |
| | CCQ410R | GCCTGAACGGGATCTTGTAGAC |
| | CCQ411F | CGGCTGTACCTCACTACGACAAAG |
| | CCQ412R | GCAAAGCCACACGCACAGTA |
| CDK1d | CCQ397F | GGCATTCCGCCTACCAGTC |
| | CCQ398R | CTCATCGCCAAGCTCCTATCG |
| CDK1e | CCQ405F | GAATGACCAAAGCAAAGATGGC |
| | CCQ406R | TGGCGTCCCAAGCACTCTG |
| | CCQ407F | CGGATCGCTTTGATATACCTTCC |
| | CCQ408R | GCCTAATATGTGATGTCAGTGATAGAAG |
| CDK2 | CCQ443F | TATGAGCCATCTCGGCGAATACCA |
| | CCQ444R | AGTTAGGCAGAACCTGTGCTCGAT |
| **Reference targets** | | |
| RPL23 | CCQ39F | CGACAGTCCAAGGCTTTCCGAAGA |
| | CCQ40R | CAGAGCCCTTCATTTCTCCCTTGT |
| EF1 | CCQ43F | AGGTCATCCCTGAACTTAACGGCA |
| | CCQ44R | GGCAGATTTGATGGCAGCGTTGAT |

#### SUPPLEMENTARY FIGURES

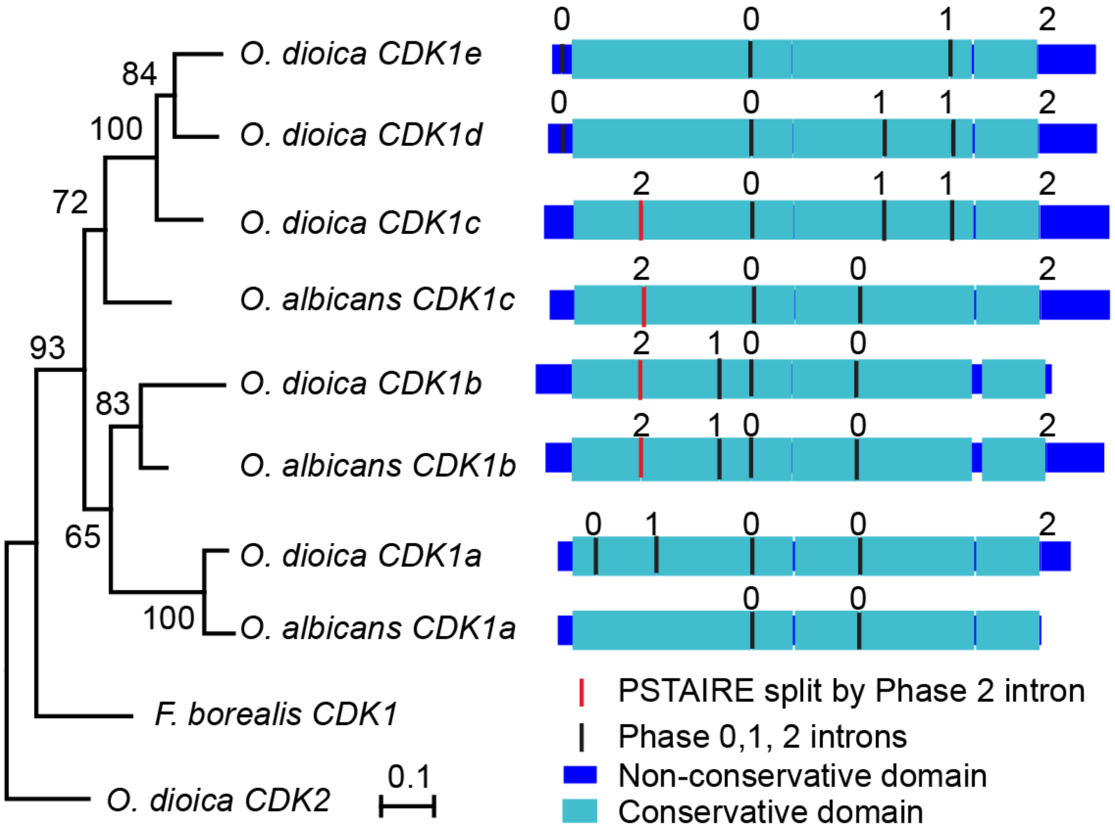


**Figure S1. Intron-exon structures of *O. dioica* and *O. albicans* CDK1 genes.** Maximum likelihood inference of selected chordate CDK1 proteins using CDK1 kinase domains with posterior probabilities indicated at nodes. Intron phases in coding regions are indicated as 0, 1 and 2. Conserved regions within *O. dioica* and *O. albicans* CDK1 sequences are indicated as light blue bars. The region encoding the PSTAIRE helix has been split by a phase 2 intron in orthologs of *Oikopleura CDK1b* and *c*.

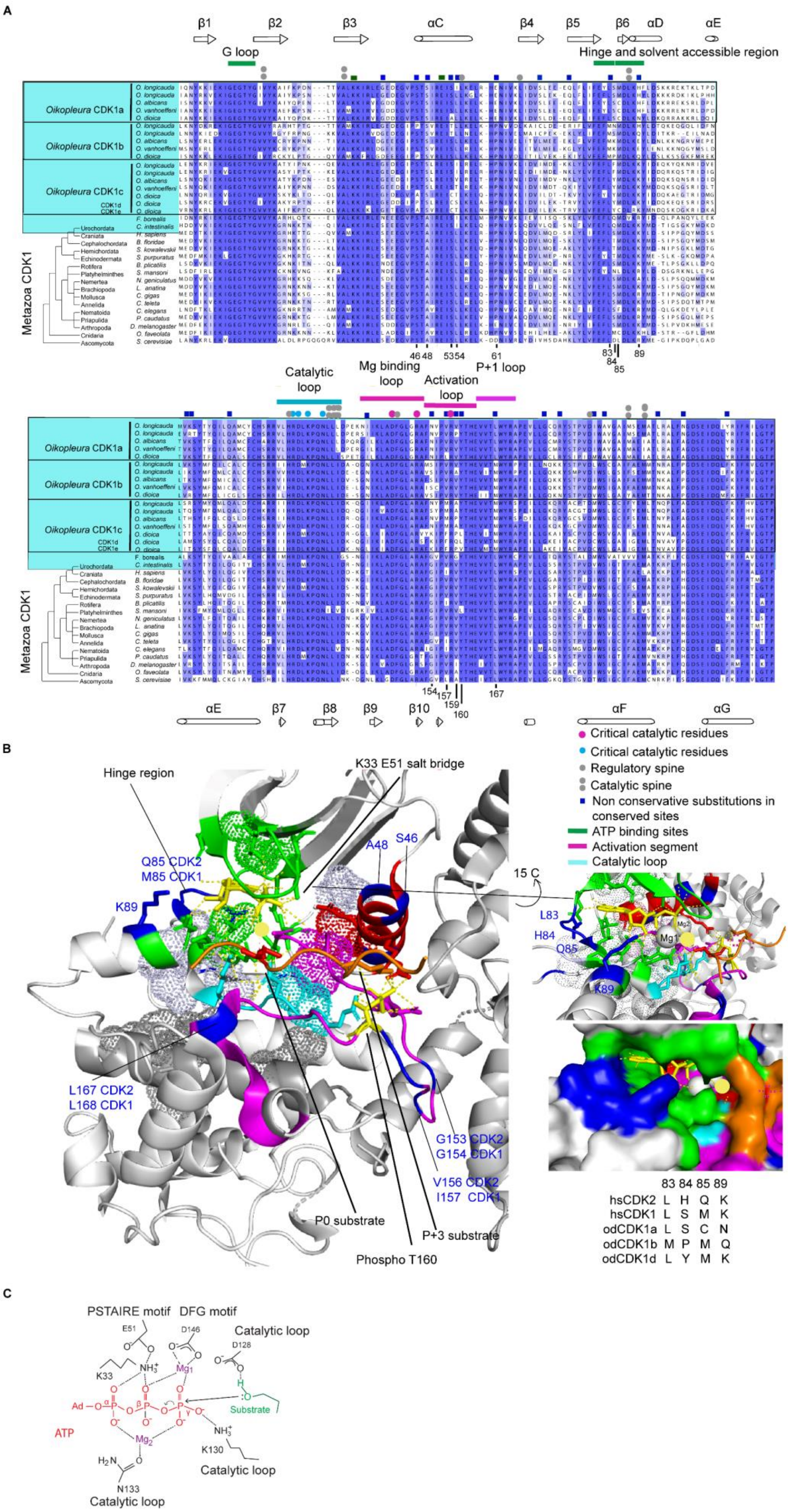

A
β1
β2
β3
αC
β4
β5
β6
αD
αE
G loop
Hinge and solvent accessible region
Oikopleura CDK1a
Oikopleura CDK1b
Oikopleura CDK1c
CDK1d
CDK1e
Metazoa CDK1
Urochordata
Craniata
Cephalochordata
Hemichordata
Echinodermata
Rotifera
Platyhelminthes
Nemertea
Brachiopoda
Mollusca
Annelida
Nematoida
Priapulida
Arthropoda
Cnidaria
Ascomycota
O. longicauda
O. albicans
O. vanhoeffeni
O. dioica
F. borealis
C. intestinalis
H. sapiens
B. floridae
S. kowalevskii
S. purpuratus
B. plicatilis
S. mansoni
N. geniculatus
L. anatina
C. gigas
C. teleta
C. elegans
P. caudatus
D. melanogaster
O. faveolata
S. cerevisiae
46 48
53 54
61
83
84
85
89
Catalytic loop
Mg binding loop
Activation loop
P+1 loop
154
157
159
160
167
β7
β8
β9
β10
αF
αG
B
Hinge region
K33 E51 salt bridge
Q85 CDK2
M85 CDK1
K89
A48
S46
L167 CDK2
L168 CDK1
G153 CDK2
G154 CDK1
V156 CDK2
I157 CDK1
P0 substrate
P+3 substrate
Phospho T160
15 C
L83
H84
Q85
K89
Mg2
Mg1
Critical catalytic residues
Critical catalytic residues
Regulatory spine
Catalytic spine
Non conservative substitutions in conserved sites
ATP binding sites
Activation segment
Catalytic loop
83 84 85 89
hsCDK2 L H Q K
hsCDK1 L S M K
odCDK1a L S C N
odCDK1b M P M Q
odCDK1d L Y M K
C
PSTAIRE motif
DFG motif
Catalytic loop
E51
D146
D128
K33
Mg1
Ad
ATP
Substrate
Mg2
K130
Catalytic loop
N133
Catalytic loop

**Figure S2. Substitutions in odCDK1 paralogs at evolutionarily conserved metazoan CDK1 residues.** (*A*) Conserved metazoan CDK1 residues identified by alignment of CDK1 sequences from 15 metazoan phyla. The budding yeast cdc28 ortholog is also included in the alignment. Substitutions in *Oikopleura* CDK1 sequences that occur at sites conserved in other metazoan CDK1s were labeled with blue squares and mapped to structural elements of the active CDK2-cyclin A1 complex crystalized with ADP, substrate peptide and $MgF_3^-$, a mimic for the γ-phosphate of ATP in the transient state (PDB ID: 3QHR). (*B*) Protein kinase structural elements are shown in white and the residues that form the regulatory and catalytic spine are shown as space-filling dots. The upper right panel shows a magnified ATP binding region and the lower panel shows a zoomed-in view of the corresponding surface-filled model. Numbering is according to human CDK2. Functional motifs are differently coloured. ADP is shown as stick model with $MgF_3^-$, a mimic for the γ phosphate group indicated as a yellow sphere in all panels. $Mg^{2+}$(1) and $Mg^{2+}$(2) are shown as white spheres in the upper right panel. Residues critical for catalysis are conserved in *Oikopleura* CDK1 paralogs including: the $Mg^{2+}$-binding loop, the catalytic loop, and the R- and C-spines. The residues associated with T160 (R50, R126, R150) are also conserved. The residues in *Oikopleura* CDK1s (adenine region, sugar region, phosphate binding region) that interact directly with ATP are conserved, whereas residues that enclose and modulate the affinity to ATP (hinge and solvent accessible region) are diversified. None of the CDK1 outparalogs have the same ATP binding sites. Again, the residues in the activation loop that direct contact with P0 and P+3 are conserved in *Oikopleura* CDK1 paralogs, but residues that modulate the conformation of the activation loop are diversified in *Oikopleura* CDK1 outparalogs. None of the *Oikopleura* CDK1 outparalogs has the same activation loop sequence. Bottom right: residues in the hinge and solvent accessible region of human (hs) CDK2, CDK1, and 3 *O. dioica* (od) CDK1 paralogs are compared. (*C*) Schematic diagram of the protein kinase catalytic mechanism. The reaction proceeds from the enzyme/substrate complex through the transition state (center) to the enzyme/product complex. Residue numbers are based on human CDK2. The OH group of the protein substrate is aligned so that the electron pair on the oxygen are in-line with the γ-phosphate (arrow) and the βγ-bridging oxygen (curved arrow) of the bound ATP.

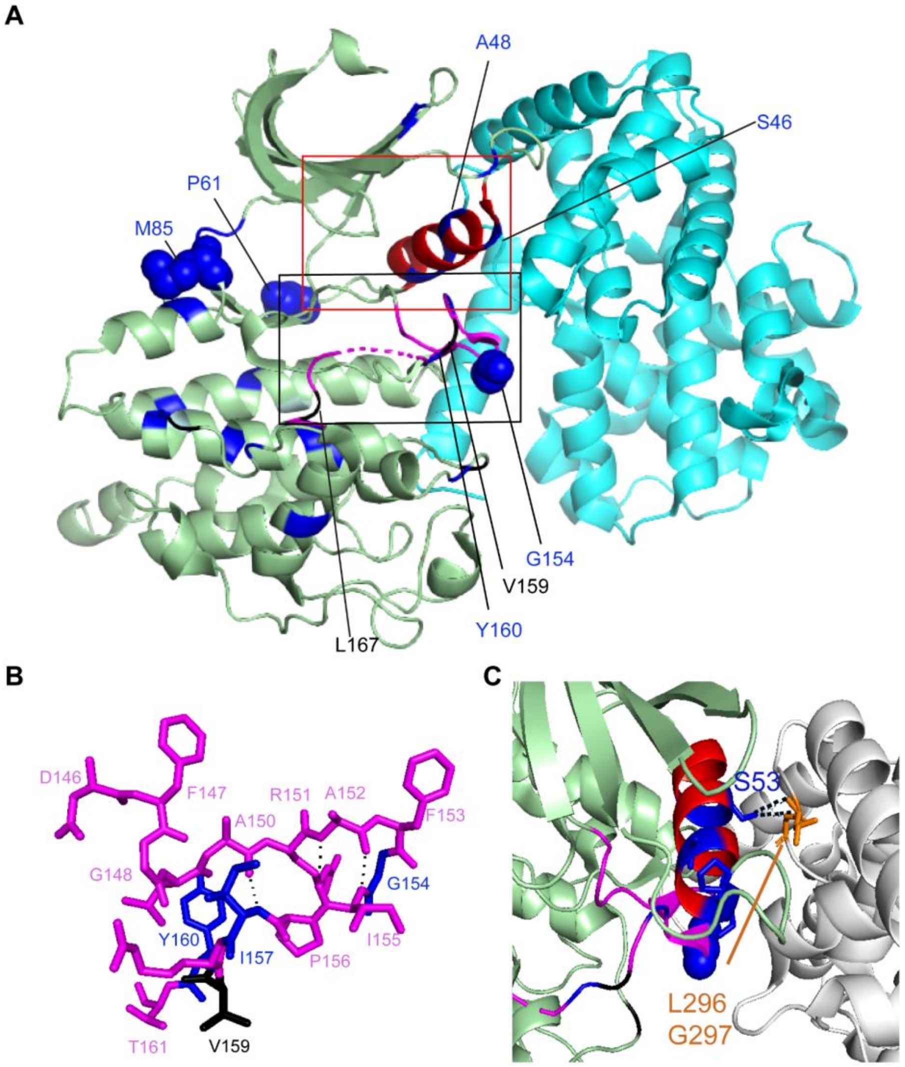


**Figure S3. Mapping of *Oikopleura* CDK1 substitution sites to the human CDK1 (green)-cyclin B1 (blue) structure, indicating positive selection sites and conserved determinants of specificity.** (*A*) Substitutions in *Oikopleura* CDK1 sequences that occur at sites conserved in other metazoan CDK1s are indicated in blue. Evolutionarily conserved determinants of substrate specificity that are modified in *Oikopleura* CDK1 paralogs are indicated in black. The conserved PSTAIRE helix, which is modified in *Oikopleura*, is indicated in red. Blue spheres indicate residues that have undergone positive selection as identified by PAML 4.1. The black box outlines the T-loop that is enlarged in (*B*). The red box outlines the PSTAIRE helix and its interface with Cyclin B that is enlarged in (*C*). (*B*) CDK1-specific β turn formed at the tip of the activation segment when CDK1 binds cyclin B1. Residues involved in the β turn are A152, F153, G154, I155 (numbering based on human CDK1). Hydrogen bonds formed between main chains are labeled with black dashed lines. **c**, CDK1-specific hydrogen bonds formed between the side chain of S53 in the PSTAIRE helix and residues L296, G297 of cyclin B1. Colour coding in (*B*) and (*C*) as in (*A*).

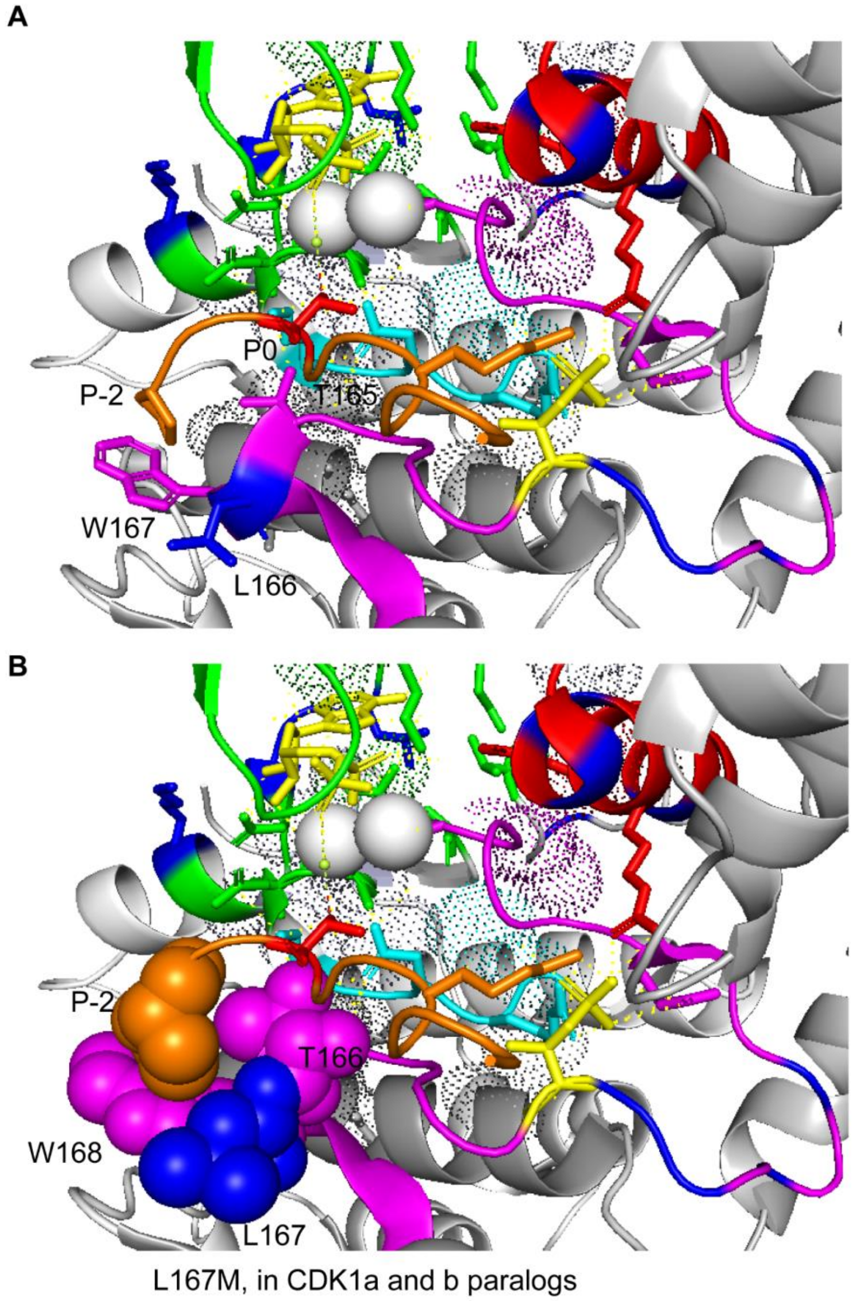


**Figure S4. CDK1/2 L167/L166 contribute to affinity toward the substrate P-2 residue.** (*A*) Active human CDK2-Cyclin A1 complex crystalized with ADP, $MgF_3^-$, a mimic for the γ-phosphate of ATP in the transient state (PDB ID: 3QHR) and substrate peptide. Residues in the activation segment of CDK2 that contact the substrate P-2 residue are coloured in magenta. or (*B*) magentas spheres. The residue numbering corresponds to the human CDK2 sequence in (*A*) and the human CDK1 sequence in (*B*). *Oikopelura* CDK1a and b paralogs share an L167M substitution. *Oikopleura* substitutions in conserved metazoan CDK1 sites are labeled in blue, the activation segment in magenta, the substrate in orange, the catalytic loop in cyan, and the PSTAIRE helix in red.

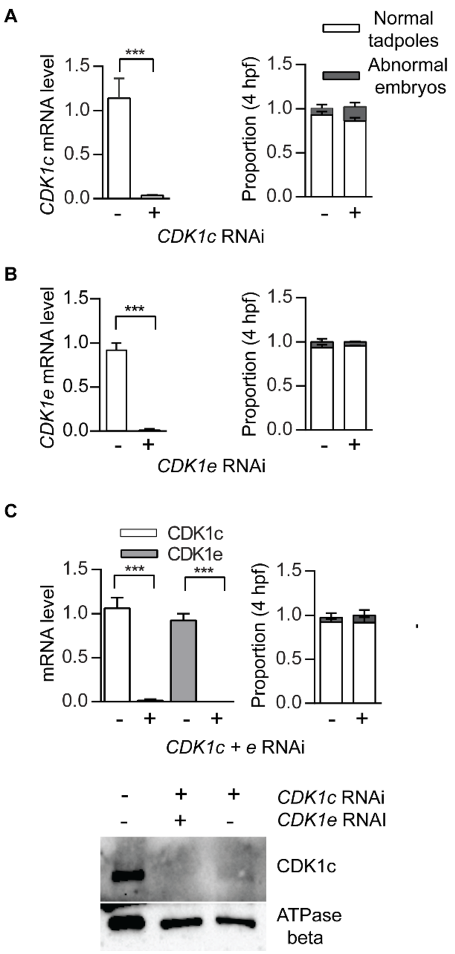


**Figure S5. CDK1c and e are dispensable for *Oikopleura* embryogenesis.** (*A*) Left panel: significant knockdown *CDK1*c in oocytes spawned from ovaries that had been injected with dsRNA against *odCDK1c* at D4. Right panel: no significant difference in developmental outcome between *CDK1c* deficient and wild type zygotes. The top legend for this panel also applies to the right panels in (*B*) and (*C*). (*B*) Left panel: significant knockdown of *CDK1e* in oocytes spawned from ovaries that had been injected with dsRNA against *odCDK1e* at D4. Right panel: no significant difference in developmental outcome between *CDK1e* deficient and wild type zygotes. (*C*) Upper left panel: significant) knockdown *CDK1c and CDK1e* in oocytes spawned from ovaries that had been injected with dsRNA against *odCDK1c* and *CDK1e* at D4 (upper left panel). Upper right panel: no significant difference in developmental outcome between *CDK1c* + *CDK1e* deficient and wild type zygotes (upper right panel). Where indicated, ***P<0.001, **P = <0.01, *P<0.05. Bottom panel: Western blot showing knockdown of CDK1c in *CDK1c* RNAi and *CDK1c* + *e* RNAi oocytes.

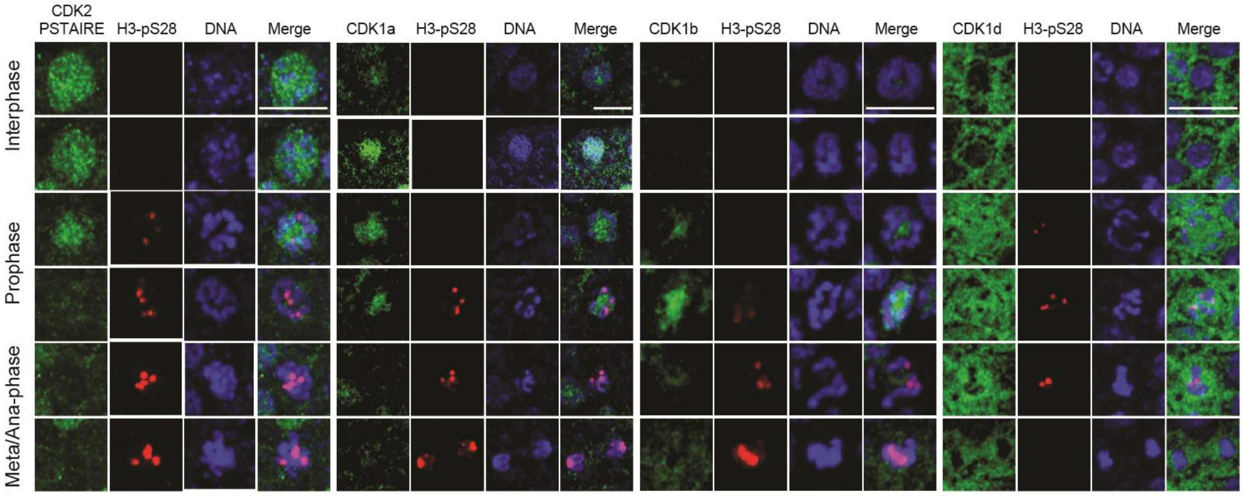


**Figure S6. Cellular localization of CDK1 paralogs during interphase and M-phase.** Antibodies were against the PSTAIRE helix of CDK2 or anti-eGFP antibodies targeting odCDK1a, b or d-eGFP fusion constructs. CDK2 disappeared on chromatin during early prophase, CDK1a initiated on chromatin during interphase and disappeared prior to metaphase. CDK1b was absent from chromatin during interphase and became intense on chromatin during the prophase to pro-metaphase, before declining on metaphase chromatin. CDK1d exhibited exclusion from chromatin until prophase when there was diffuse staining on chromatin, which largely disappeared in metaphase. Scale bars: 5 µm.

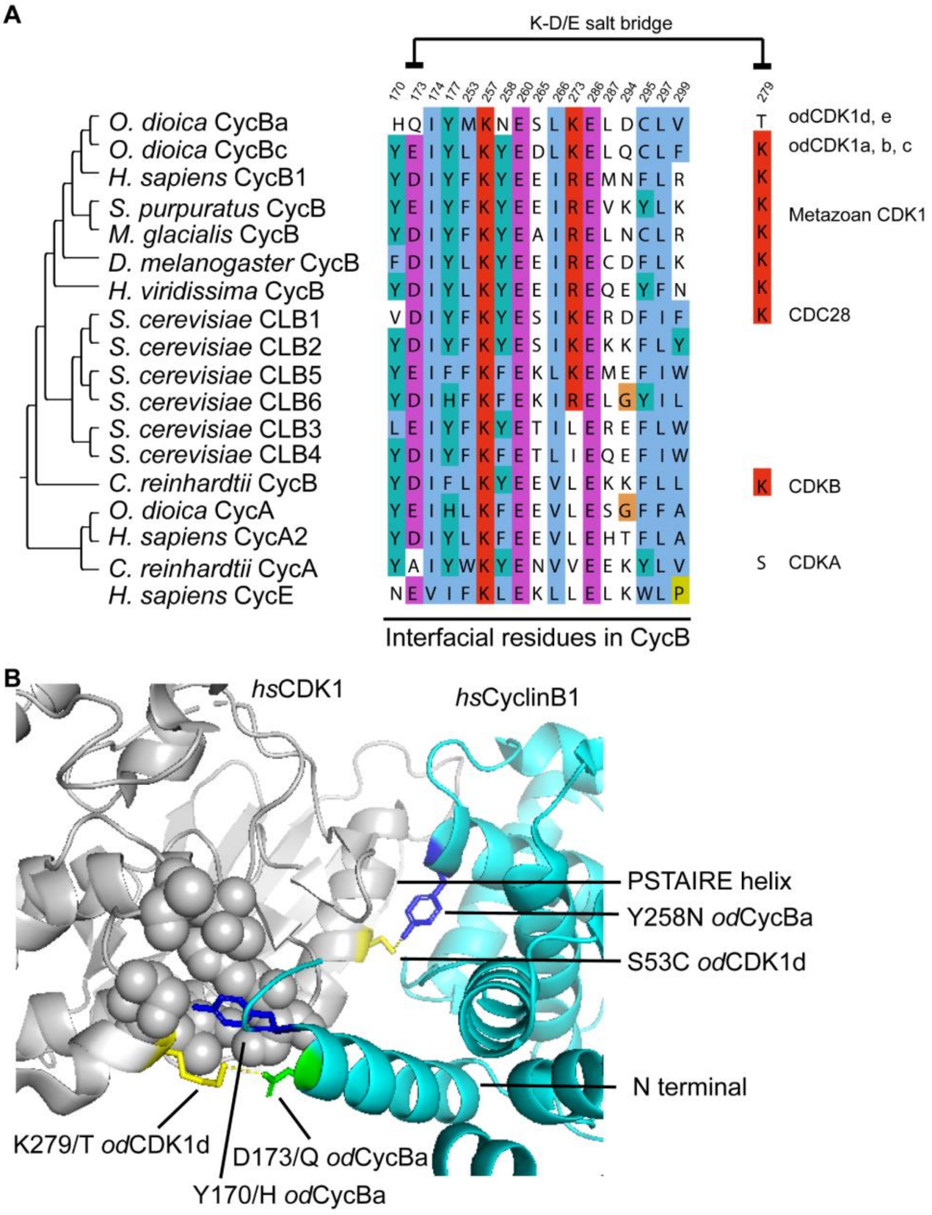


**Figure S7. *Oikopleura* CDK1d:Cyclin Ba-specific substitutions in the interfacial region.** (*A*) Alignment of amino acids in Cyclin B homologs that mediate the CDK1:Cyclin B interface, as identified using CCP4 CONTACT (PDB ID: 4YC3). (*B*) Human CDK1:Cyclin B1 structure (PDB ID: 4YC3) showing conserved contacts between the Cyclin B1 N-terminal Y170 and the CDK1 C-lobe hydrophobic pocket. Cyclin B substitutions at the Y170 site were present in odCyclin Ba, not in Bc. The salt bridge formed between CycB1 E173 and CDK1 K279 is conserved in metazoans and plants, but this pair of residues has been modified in odCDK1d (or e):CycBa pairings. Sequences were from *Oikopleura dioica*, *Homo sapiens*, *Strongylocentrotus purpuratus*, *Marthasterias glacialis*, *Drosophila melanogaster*, *Hydra viridissima*, *Saccharomyces cerevisiae*, and *Chlamydomonas reinhardtii*.
